# Epigenetic control of Topoisomerase 1 activity presents a cancer vulnerability

**DOI:** 10.1101/2024.10.22.619113

**Authors:** Tae-Hee Lee, Colina X Qiao, Vladislav Kuzin, Yuepeng Shi, Vijayalalitha Ramanaranayan, Tongyu Wu, Xianzhen Zhou, David Corujo, Marcus Buschbeck, Laura Baranello, Philipp Oberdoerffer

**Author notes:** Lead Contact: 1550 Orleans St, Baltimore MD 21287.

## Abstract

DNA transactions introduce torsional constraints that pose an inherent risk to genome integrity. While topoisomerase 1 (TOP1) activity is essential for removing DNA supercoiling, aberrant stabilization of TOP1:DNA cleavage complexes (TOP1ccs) can result in cytotoxic DNA lesions. What protects genomic hot spots of topological stress from aberrant TOP1 activity remains unknown. Here, we identify chromatin context as an essential means to coordinate TOP1cc resolution. Through its ability to bind poly(ADP-ribose) (PAR), a protein modification required for TOP1cc repair, the histone variant macroH2A1.1 establishes a TOP1-permissive chromatin environment, while the alternatively spliced macroH2A1.2 isoform is unable to bind PAR or protect from TOP1ccs. By visualizing transcription-induced topological stress in single cells, we find that macroH2A1.1 facilitates PAR-dependent recruitment of the TOP1cc repair effector XRCC1 to protect from ssDNA damage. Impaired macroH2A1.1 splicing, a frequent cancer feature, was predictive of increased sensitivity to TOP1 poisons in a pharmaco-genomic screen in breast cancer cells, and macroH2A1.1 inactivation mirrored this effect. Consistent with this, low macroH2A1.1 expression correlated with improved survival in cancer patients treated with TOP1 inhibitors. We propose that macroH2A1 alternative splicing serves as an epigenetic modulator of TOP1-associated genome maintenance and a potential cancer vulnerability.

## Introduction

Single-stranded DNA (ssDNA) lesions (SSLs) are among the most abundant DNA aberrations and pose a pervasive threat to genome integrity ^1^. SSLs frequently arise during transcription- or DNA replication-associated topological stress as a result of DNA topoisomerase 1 (TOP1)-mediated DNA supercoil resolution ^2^. TOP1 unwinds DNA by nicking one strand of the double-helix, creating a covalent TOP1:DNA cleavage complex (TOP1cc) and concomitant ssDNA break as transient intermediates ^3^. The TOP1 catalytic cycle can be aborted in response to reactive metabolites, aberrant repair in the presence of adjacent base damage or non-B DNA structures, as well as in response to anticancer drugs ^3^. Unresolved TOP1ccs have been linked to somatic mutations, chromosomal aberrations and cell death. Consistent with this, genetic defects in TOP1cc repair can cause human genome instability syndromes, and TOP1cc stabilization is the mechanism of action of various anticancer drugs ^2,4–6^. Tight regulation of topoisomerase function is thus essential to ensure genome integrity, and its manipulation presents a potential cancer vulnerability.

Chromatin composition has emerged as an effective mediator of genome maintenance ^7,8^. However, we know surprisingly little about how chromatin affects the repair of SSLs. Topoisomerase enzymes are often concentrated in genomic regions prone to recurrent topological stress, emphasizing the need to promptly resolve torsional constraints before they disrupt the underlying genetic processes ^2^. What directs TOP1 function to hotspots of DNA supercoiling while simultaneously preventing excessive TOP1cc accumulation at these regions is a central, albeit unresolved, question in topological genome maintenance.

TOP1cc resolution involves the activation of Poly(ADP-ribose) (PAR) Polymerase 1 (PARP1), which senses ssDNA breaks that occur during the TOP1 catalytic cycle and orchestrates TOP1cc degradation, ssDNA gap filling and DNA re-ligation through PARylation of TOP1ccs and downstream repair effectors ^1,9^. How PAR-dependent repair events are coordinated at the site of damage remains unknown. MacroH2A1.1, one of two alternatively spliced isoforms of the macro-histone variant macroH2A1, is the only nucleosome component with an inherent ability to bind PAR ^10–12^. Consistent with a role in the orchestration of PARP-driven DNA repair processes, we previously discovered that macroH2A1.1, but not the PAR-binding-deficient macroH2A1.2 splice isoform, acts as an effector of microhomology-mediated end joining (MMEJ) through PARP-dependent interaction with MMEJ repair factors ^13^. Moreover, macroH2A1.1 was found to accumulate at the TSS of active genes ^14–16^, a hotspot of TOP1 binding and PARP1 activity, which is nevertheless protected from TOP1cc accumulation^17,18^.

Here we identify macroH2A1.1 as an epigenetic rheostat of TOP1 function that protects from TOP1-dependent DNA lesions by coordinating PAR-dependent TOP1cc repair. MacroH2A1.1 alternative spicing is frequently perturbed in cancer ^10^, and we find that impaired macroH2A1.1 splicing is predictive of increased sensitivity to TOP1 poisons in cancer cell lines. Conversely, high macroH2A1.1 expression correlated with decreased SSL-related mutation signatures in tumor tissue and poor survival outcome in ovarian cancer patients treated with TOP1 inhibitors (TOP1i). Two main conclusions arise from this work: (1) chromatin composition, and specifically TOP1-associated macroH2A1.1 domains (TMDs), control the response to topological stress, and (2) epigenetic changes via the manipulation of macroH2A1.1 splicing can alter TOP1cc repair outcome, presenting a cancer vulnerability. Our findings have direct implications for transcription-associated genome instability as well as cancer therapy.

## Results

### macroH2A1.1 defines a TOP1-permissive chromatin environment

Prompted by the observation that both TOP1 and macroH2A1.1 are enriched at the TSS of active genes ^15,17^, we asked whether colocalization with macroH2A1.1 is a general hallmark of chromatin-bound TOP1. To test this, we performed Cleavage Under Targets and Release Using Nuclease (CUT&RUN) followed by next generation sequencing (NGS) for macroH2A1.1 and TOP1 in MDA-MB-231 breast cancer cells, which express relatively high levels of macroH2A1.1 ^19^. To ensure isoform-specific detection of macroH2A1.1, we generated two independent macroH2A1.1 knockout (1.1-KO) clones reconstituted with FLAG-tagged macroH2A1.1 ^20^. Parental MDA-MB-231 cells or cells reconstituted with empty vector served as negative controls for FLAG-macroH2A1.1 CUT&RUN specificity (**Fig. 1A**). Replicate NGS experiments were performed for each condition (Spearman correlation coefficient r > 0.9), and good concordance was observed between the two knockout clones (r ∼ 0.7, **Fig. S1A**). Consistent with previous reports, macroH2A1.1 peaks were highly correlated with heterochromatin domains marked by K27-trimethylated histone H3 (H3K27m3), validating our FLAG-IP approach (**Fig. 1B, C, Fig S1B**) ^14,15,21^. We then compared macroH2A1.1 and TOP1 chromatin profiles, which revealed robust colocalization of the two proteins at TSS-proximal TOP1 peaks (**Fig. 1A, B**). Remarkably, macroH2A1.1/TOP1 colocalization was not restricted to the TSS but extended to most if not all TOP1-enriched genomic regions, pointing to a general affinity of macroH2A1.1 for sites of TOP1 function (**Fig. 1A**). Non-random colocalization of macroH2A1.1 and TOP1 was confirmed using the Jaccard index, a statistic for measuring sample set similarity between two peak set by comparing the overlap between observed and randomly shuffled (permuted) peaks of equal size (**Fig. 1C**). The overlap of macroH2A1.1 at TOP1 peaks was comparable to that of PARP1, a known interactor of both TOP1 and macroH2A1.1 ^14,22,23^. TOP1-associated macroH2A1.1 domains (TMDs) were notably distinct from the well-characterized, heterochromatin-associated macroH2A1.1 domains, as they were depleted for H3K27me3 (**Fig. 1C**). We thus propose that TMDs present a unique macroH2A1.1 chromatin environment reminiscent of macroH2A1 regions previously associated with PARP-dependent gene regulation ^14,15,24^.

**Figure 1.**
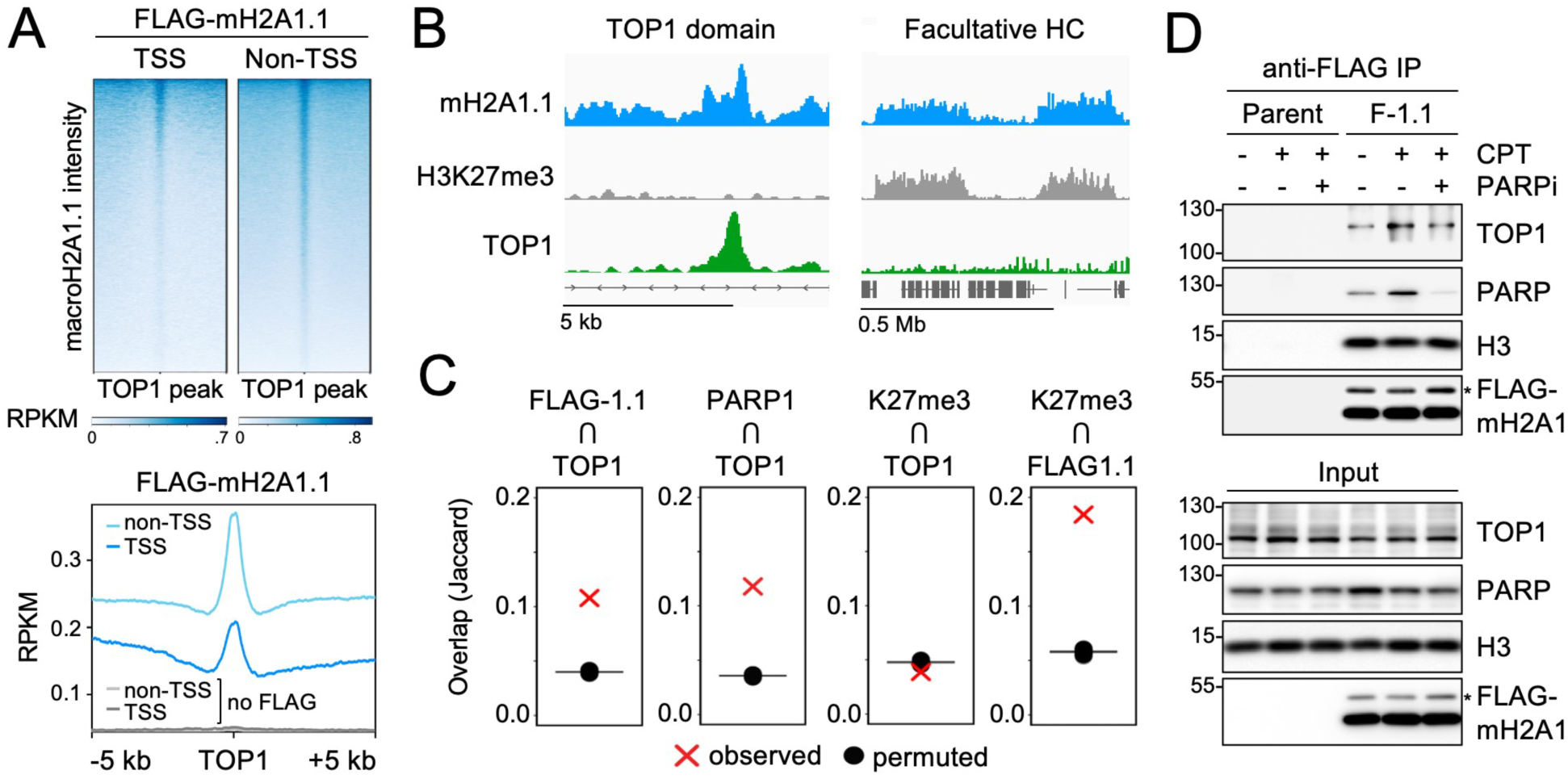
MacoH2A1.1 defines TOP1 permissive chromatin domains. **(A)** Heatmaps and profile plots for FLAG-macroH2A1.1 CUT&RUN signal centered on TSS-proximal (TSS) and TSS-distal (non-TSS) TOP1 peaks in MDA-MB-231 macroH2A1.1 KO cells reconstituted with FLAG-macroH2A1.1. Cells without FLAG-macroH2A1.1 served as negative control (no FLAG), a representative CUT&RUN experiment is shown. **(B)** IGV browser shots of distinct macroH2A1.1 chromatin environments associated with TOP1 domains (left) or facultative heterochromatin (HC) marked by H3K27me3 (right). **(C)** Jaccard indices for observed peak overlap between a feature of interest (top) and a reference feature (bottom), or randomly shuffled reference feature peaks of equal size (permuted). Values on the y-axis represent the intersection divided by the union in base pairs. **(D)** Western blot for the indicated proteins in nuclear lysates (input) or anti-FLAG IP samples from parental and FLAG-macroH2A1.1 knock-in 293 cells (F-1.1) in the presence or absence of CPT and the PARPi Olaparib. * ubiquitinated macroH2A1.

To assess if macroH2A1.1 colocalization with TOP1 involves physical association between the two proteins, we performed co-immunoprecipitation (Co-IP) in 293 cells expressing FLAG-tagged macroH2A1.1. We observed robust interaction between TOP1 and macroH2A1.1 that increased 2 to 3-fold upon treatment with the TOP1i camptothecin (CPT), which stabilizes catalytically engaged TOP1ccs (**Fig. 1D**). Little interaction was observed with the alternatively spliced macroH2A1.2 isoform expressed at comparable levels (**Fig. S1C**). Corroborating a role for macroH2A1.1 in the cellular response to topological stress, the CPT-induced increase in TOP1:macroH2A1.1 association was reverted to pre-damage levels in the presence of the PARP inhibitor (PARPi) Olaparib. Moreover, PARP1 itself showed a similar, PARP activity-dependent increase in macroH2A1.1 association upon CPT treatment (**Fig. 1D**). The interaction of macroH2A1.1 with core histone H3 was unaffected by either treatment, demonstrating comparable IP efficiency. We therefore conclude that macroH2A1.1 associates with TOP1 on chromatin in a splice isoform-specific manner that is enhanced upon TOP1cc damage.

### macroH2A1.1 protects from TOP1:DNA lesions

We next sought to determine if macroH2A1.1 affects TOP1cc accumulation and/or resolution. To this end, we performed TOP1 Covalent Adduct Detection sequencing (TOP1 CAD-Seq, **Fig. 2A**) ^25^, which maps genomic regions harboring catalytically engaged TOP1. Consistent with previous reports ^17^, we found TOP1ccs to be depleted from the TSS of active genes despite significant TOP1 accumulation (**Fig. 2B**), suggesting that they are efficiently cleared to ensure genome integrity at these regions of high torsional stress ^26^. To test this possibility, we devised a modified CAD-Seq approach that measures TOP1cc resolution upon damage relative to steady state TOP1cc levels. For steady state TOP1cc detection, cells were briefly pulsed with CPT to stabilize existing TOP1:DNA intermediates in the presence of the proteasome inhibitor MG132, which inhibits repair-mediated TOP1cc degradation and removal ^27^. Negligible CAD-Seq signal was observed in the absence of CPT, demonstrating assay specificity (**Fig. S2A**) ^25^. To assess TOP1cc resolution after prolonged damage, cells were treated with CPT for 30 min in the absence of MG132, causing extended TOP1cc trapping while simultaneously allowing for repair of the lesion. Replicate NGS experiments were performed for each condition and combined for downstream analyses (r ≥ 0.85, **Fig. S2B**). The turnover of damage-induced TOP1:DNA adducts was defined as the ratio of TOP1cc levels after prolonged CPT exposure over the steady state (ΔTOP1cc), and assessed in the presence or absence of stable macroH2A1.1 knockdown (**Fig. 2C, Fig. S2C**). MacroH2A1.1 depletion resulted in a significant, TSS-proximal increase in ΔTOP1cc, pointing to increased TOP1cc accumulation, or impaired TOP1cc turnover, upon damage (**Fig. 2C**). The extent of TOP1cc accumulation directly correlated with gene expression levels and was most pronounced at highly transcribed genes, where torsional stress is maximal (**Fig. 2D**) ^17,26^. In contrast, even highly expressed genes showed efficient TOP1cc turnover (low ΔTOP1cc) in the presence of macroH2A1.1, consistent with effective TOP1cc clearance. Extending our findings beyond the TSS, we observed a similar pattern of macroH2A1.1-dependent TOP1cc turnover across TOP1 peaks genome-wide, which was directly correlated with the extent of macroH2A1.1 enrichment. In other words, TOP1 peaks with the highest macroH2A1.1 colocalization showed increased CPT-induced TOP1cc accumulation upon macroH2A1.1 loss, while TOP1 peaks with low macroH2A1.1 colocalization were only marginally affected (**Fig. 2E**). Of note, macroH2A1.1 loss-associated TOP1cc accumulation extended into the flanking regions of macroH2A1.1^high^ TOP1 peaks, consistent with abundant macroH2A1.1 enrichment beyond the TOP1 peak at these sites (**Fig. 2E**). No change in overall TOP1 expression was observed upon macroH2A1.1 inactivation (**Fig. S3A**). These findings support a functional link between TOP1cc turnover and macroH2A1.1.

**Figure 2.**
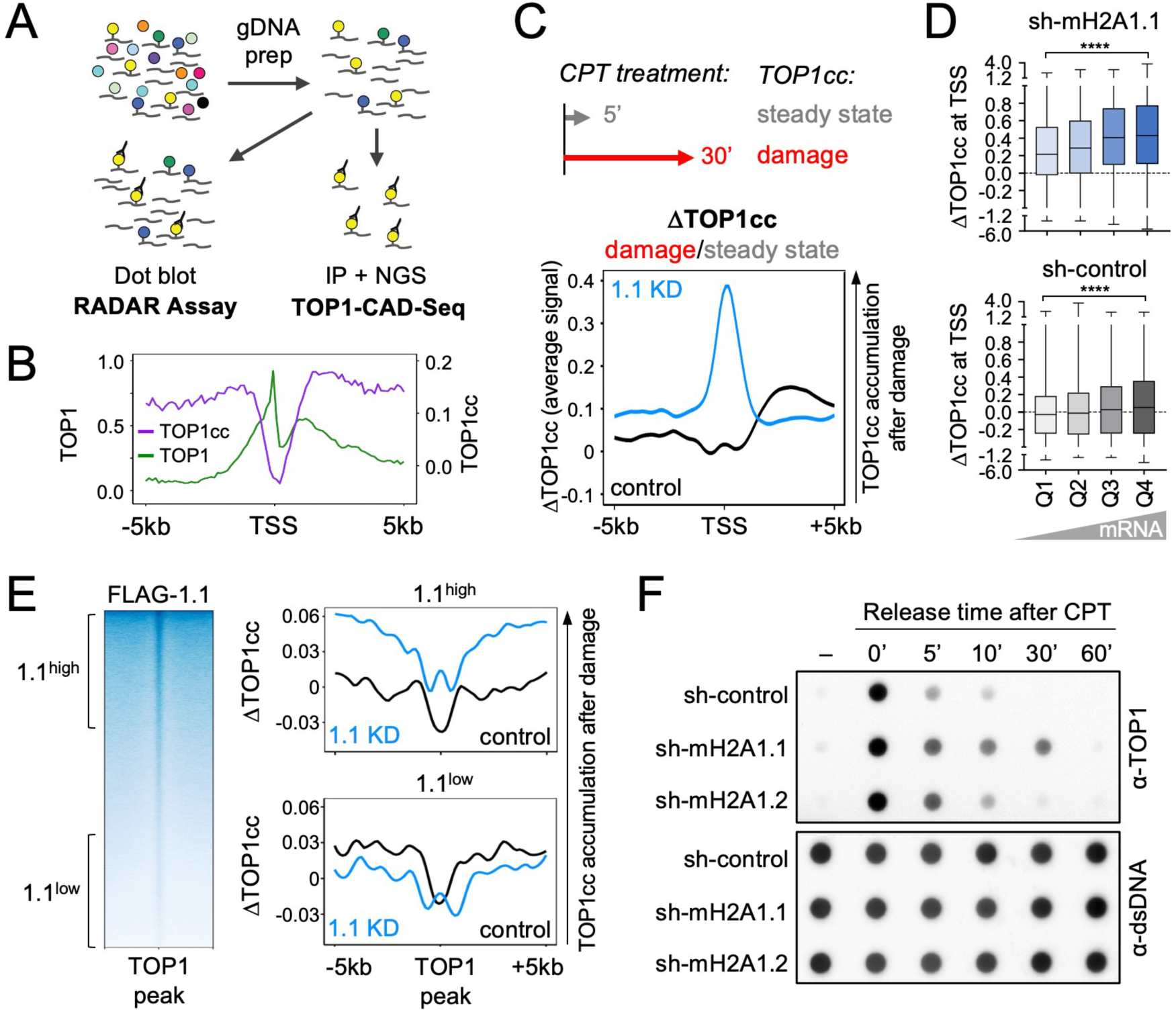
MacroH2A1.1 protects from TOP1cc accumulation. **(A)** Schematic for RADAR and TOP1 CAD-Seq assays. **(B)** Representative TSS profile plots for TOP1 CUT&RUN (green) and TOP1cc CAD-Seq (purple) for the top quartile of expressed genes in MDA-MB-231 cells, as defined by RNA-Seq ^15^. Y-axes depict Z-normalized read counts. **(C)** TSS-associated TOP1cc resolution following CPT-induced damage in the presence or absence of macroH2A1.1. Profile plot depicts Z-normalized DTOP1cc ratios of damage-induced (30’ CPT) over steady state TOP1cc (5’ CPT) for sh-RFP control (black) and macroH2A1.1 knockdown (1.1 KD, blue). **(D)** Violin plots based on samples in (C) depicting mean DTOP1cc ratios within 500 bp of the TSS, separated based on RNA-Seq-derived gene expression quartiles; Q1: bottom 25%, Q4: top 25%. **** p < 3e-16 based on Wilcoxon rank-sum t-test, all quartiles are significantly different between 1.1 KD and control shRNA (p < 3e-16). **(E)** TOP1 peak-associated DTOP1cc ratios as in (C). TOP1 peaks were separated into top (1.1^high^) and bottom tertiles (1.1^low^) based on macroH2A1.1 (FLAG-1.1) enrichment, see TOP1-centered FLAG-1.1 heatmap. **(F)** TOP1 RADAR assay with genomic DNA from MDA-MB-231 cells expressing the indicated shRNAs, prior to and at the indicated timepoints after CPT treatment (1 µM, 30 min). See Fig. S2D for a quantification of two independent experiments.

To dissect the underlying TOP1cc repair kinetics, we measured TOP1cc levels in total genomic DNA using the Rapid Approach to DNA Adduct Recovery (RADAR) ^28^, which, like TOP1 CAD-Seq, specifically detects TOP1ccs, but uses DNA dot blot instead of IP/NGS (**Fig. 2A**). RADAR was performed in cells treated with CPT for 30 min followed by drug wash-out and varying recovery times in the presence or absence of either macroH2A1 splice isoform. While macroH2A1 loss did not affect overall TOP1cc induction upon CPT treatment, TOP1cc levels remained elevated 30 min after CPT removal specifically in macroH2A1.1 depleted cells, indicative of delayed TOP1cc repair (**Fig. 2F, Fig. S2C, D**). Taken together, these findings demonstrate that macroH2A1.1 acts in an isoform-specific manner to regulate TOP1cc clearance at genomic regions of high TOP1 activity.

### PAR-binding-deficient macroH2A1.1 impairs TOP1cc clearance

The selective impact of macroH2A1.1 on TOP1cc repair prompted us to determine if this function is dependent on its ability to bind PAR. To test this, we engineered MDA-MB-231 1.1-KO cells that exclusively express a PAR-binding-deficient macroH2A1.1 mutant (macroH2A1.1 G224E ^29^) and assessed its impact on TOP1cc turnover following CPT treatment compared to 1.1-KO cells or cells reconstituted with wild-type macroH2A1.1 (**Fig. 3A**). While both macroH2A1.1 chromatin enrichment and TOP1 expression were comparable in cells expressing either wildtype or G224E mutant protein (**Fig. S1A**, **Fig. S3A, B**), the PAR-binding mutant displayed a significant delay in TOP1cc repair relative to wildtype macroH2A1.1, as did 1.1-KO cells (**Fig. 3A, B**). A similar result was observed upon overexpression of wild-type or G224E mutant macroH2A1.1-GFP fusion proteins in MCF7 breast cancer cells, which express inherently low levels of macroH2A1.1 (**Fig. S3C, Fig. 6A**) ^19^. ^19^. Together, these findings support a PAR-dependent role for macroH2A1.1 in protecting regions of topological stress from TOP1cc accumulation.

**Figure 3.**
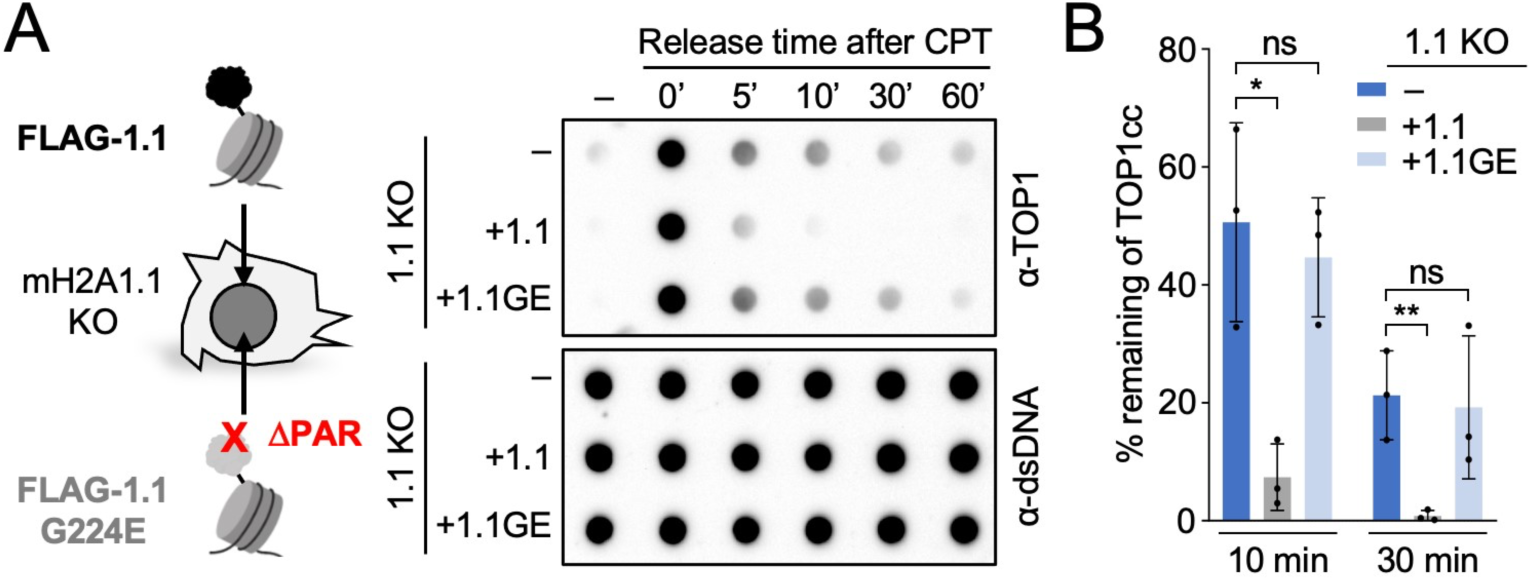
TOP1cc clearance depends on the macroH2A1.1 PAR binding domain. **(A)** TOP1 RADAR assay with genomic DNA from macroH2A1.1 knockout MDA-MB-231 cells reconstituted with WT (+1.1) or G224E mutant FLAG-macroH2A1.1 (+1.1GE), prior to and at the indicated timepoints after CPT treatment, one of three representative experiments is shown. (**B**) Quantification of RADAR analysis in (A), depicting the percentage of remaining TOP1cc relative to 0’ after CPT treatment (n = 3), values are expressed as mean and SD. P values are based on student’s two-tailed t-test, * p < 0.05, ** p < 0.01.

### macroH2A1.1 links PARP1 activity to TOP1 repair factor assembly

Having identified macroH2A1.1 as an epigenetic effector of TOP1cc turnover, we sought to determine how this histone variant modulates the resolution of trapped TOP1ccs. We have previously characterized macroH2A1 isoform interactomes using differential metabolic labeling of 293 cells expressing FLAG-tagged macroH2A1.1 or macroH2A1.2 protein ^30^. Supporting a splice isoform-specific role for macroH2A1.1 in response to single-stranded DNA lesions, three out of five macroH2A1.1 interactors, LIG3, XRCC1, and PARP1, represent repair factors that are involved in base excision repair (BER), a pathway required for the resolution of TOP1cc-associated DNA lesions ^13^. Given that XRCC1 depends on PAR/PARP1 for efficient recruitment to sites of damage ^31,32^, we asked if its association with macroH2A1.1 affects XRCC1 function in TOP1cc repair. Using Co-IP, we confirmed that XRCC1 and macroH2A1.1 interact in a PARP activity-dependent manner, while little interaction was observed with the PAR binding-deficient macroH2A1.2 isoform (**Fig. 4A**). To determine functional relevance for TOP1cc repair, we assessed XRCC1 recruitment to CPT-induced DNA lesions in the presence or absence of macroH2A1.1, using immunofluorescence (IF) imaging in MCF7 cells. CPT treatment caused a significant increase in nuclear XRCC1 foci, as reported previously ^33^, which was blunted in macroH2A1.1-deficient cells and fully restored by wild-type macroH2A1.1 but not the G224E mutant (**Fig. 4B-D, Fig. S4A, B**). Consistent with this, we observed a CPT-induced increase in Co-IP of XRCC1 with FLAG-macroH2A1.1, which was dependent on both the macroH2A1.1 PAR binding domain and PARP1 activity (**Fig. S4D**). Cell cycle profiles and overall XRCC1 expression were comparable across cell lines and experimental conditions (**Fig. S4B, C**). Moreover, defects in XRCC1 foci formation were apparent in non-S phase cells, demonstrating that this observation does not dependent on replication-associated topological stress (**Fig. S4E**) ^34^. Finally, macroH2A1.1-depleted cells showed a significant increase in CPT-induced ssDNA lesions, consistent with impaired TOP1cc-associated ssDNA break resolution (**Fig. 4E**). These findings demonstrate a role for macroH2A1.1 in XRCC1 recruitment and SSL repair that is at least in part dependent on its ability to bind PAR, establishing macroH2A1.1 as an effector of BER repair factor assembly downstream of PARP.

**Figure 4.**
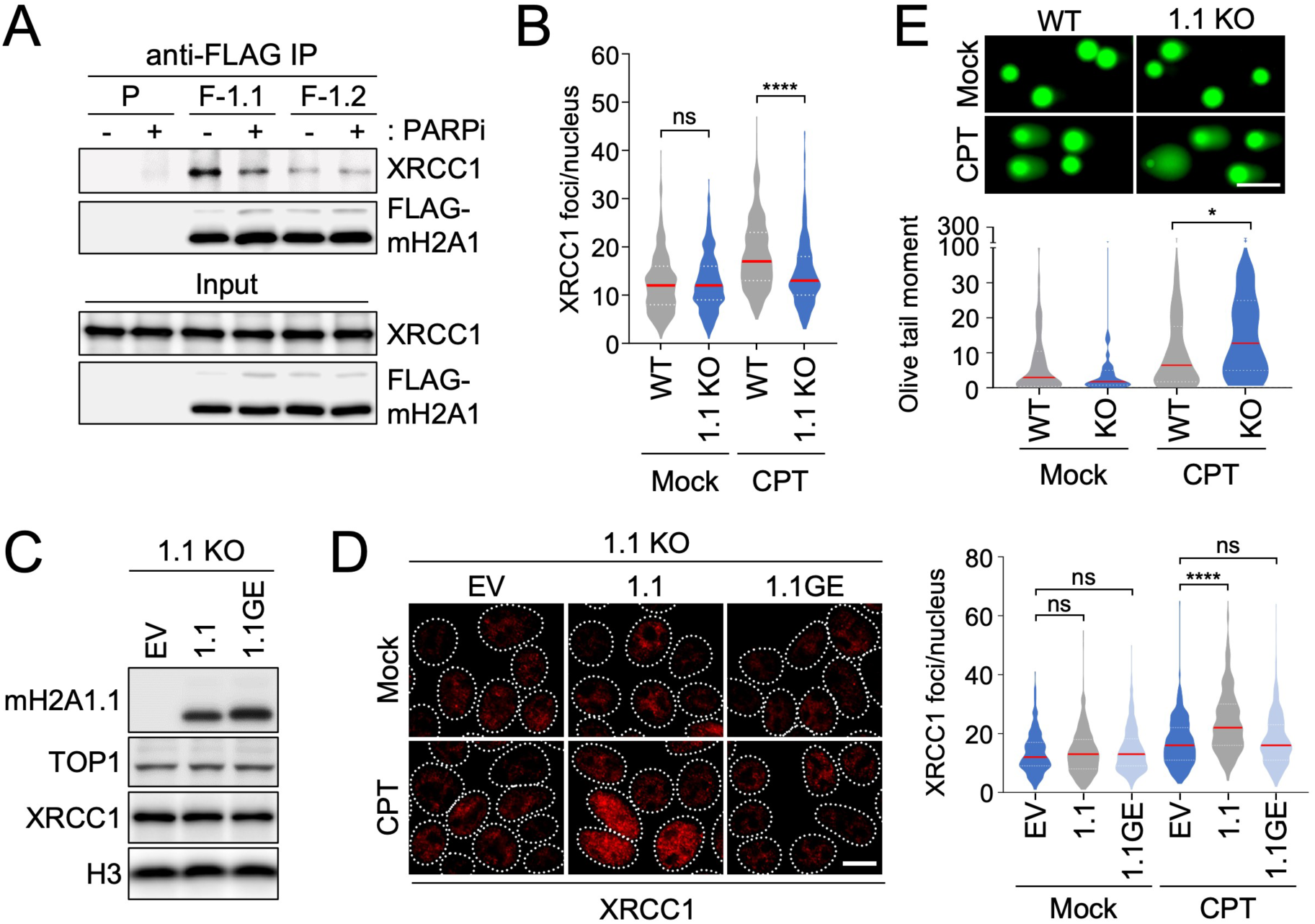
XRCC1 recruitment to TOP1ccs depends on macroH2A1.1. **(A)** Western blot for the indicated proteins in nuclear lysates (input) or IP lysates from parental (P) and FLAG-macroH2A1.1 (F-1.1) or FLAG-macroH2A1.2 (F-1.2) knock-in 293 cells in the presence or absence of PARPi. **(B)** Quantification of XRCC1 foci in MCF7 WT and macroH2A1.1 KO cells in the presence or absence of CPT treatment (1 µM, 30 min), y-axis depicts foci per nucleus (n>350). Representative images are shown in Fig. S4A. **(C)** Western blot for the indicated proteins in macroH2A1.1 KO (1.1 KO) MCF7 cells reconstituted with empty vector (EV), FLAG-macroH2A1.1 (1.1) or FLAG-macroH2A1.1 G224E (1.1GE). **(D)** XRCC1 foci in cells from (C) treated with CPT (1 µM, 30 min) or DMSO (Mock). One of two independent experiments is shown, scale bar: 10 µm. Cells were quantified as in (B), n>300 nuclei per sample. A representative of two independent clonal 1.1 KO reconstitutions is shown. **(E)** Alkaline comet assay in WT and 1.1 KO MCF7 cells treated with 20 µM CPT or DMSO (Mock); y-axis depicts Olive tail moment (n>50), a representative of two independent experiments is shown, scale bar: 100 µm. For all violin plots, red lines reflect the median, P values are based on Mann-Whitney U test; ** p < 0.01, **** p < 0.0001, ns: not significant.

### macroH2A1.1 promotes SSL repair at sites of nascent transcription

Our genome-wide data point to the TSS as a hotspot of macroH2A1.1-mediated TOP1cc repair. We thus asked if macroH2A1.1 repair function can be monitored specifically at transcription-induced DNA lesions. In contrast to DSBs, no tools exist to our knowledge for the image-based detection of locally defined TOP1cc repair events. To overcome this limitation, we have adapted a previously described transcriptional reporter system to visualize the accumulation and turnover of TOP1cc repair factors at an actively transcribed genomic locus in U2OS cells ^35,36^. Following reporter gene induction with doxycycline (Dox), the site of transcription was detected via the accumulation of yellow fluorescent protein (YFP)-tagged viral MS2 coat protein (YFP-MCP) bound to 24 MS2 stem-loop repeats within the nascent transcript (**Fig. 5A**). As a readout for TOP1 activity and associated repair events, we assessed the accumulation of TOP1 and XRCC1 based on average IF signal intensities across at least 50 MS2 sites. An MS2-distal region of equal size was analyzed in parallel to control for background signal intensity (**Fig. 5A**). To prevent PAR chain degradation and thereby stabilize repair factor accumulation upon MS2 induction, cells were treated for 30 min with an inhibitor of poly(ADP-ribose) glycohydrolase inhibitor (PARGi) ^37^. Consistent with previous reports, we observed robust MS2 signal 5 h after Dox treatment ^35,36^. MS2 induction was accompanied by an accumulation of both XRCC1 and TOP1 at and/or adjacent to the MS2 signal, indicative of transcription-associated engagement of the BER pathway even in the absence of CPT-induced TOP1cc stabilization (**Fig. 5B**). No focal XRCC1 or TOP1 enrichment was observed at the MS2-distal control region (**Fig. 5B**).

**Figure 5.**
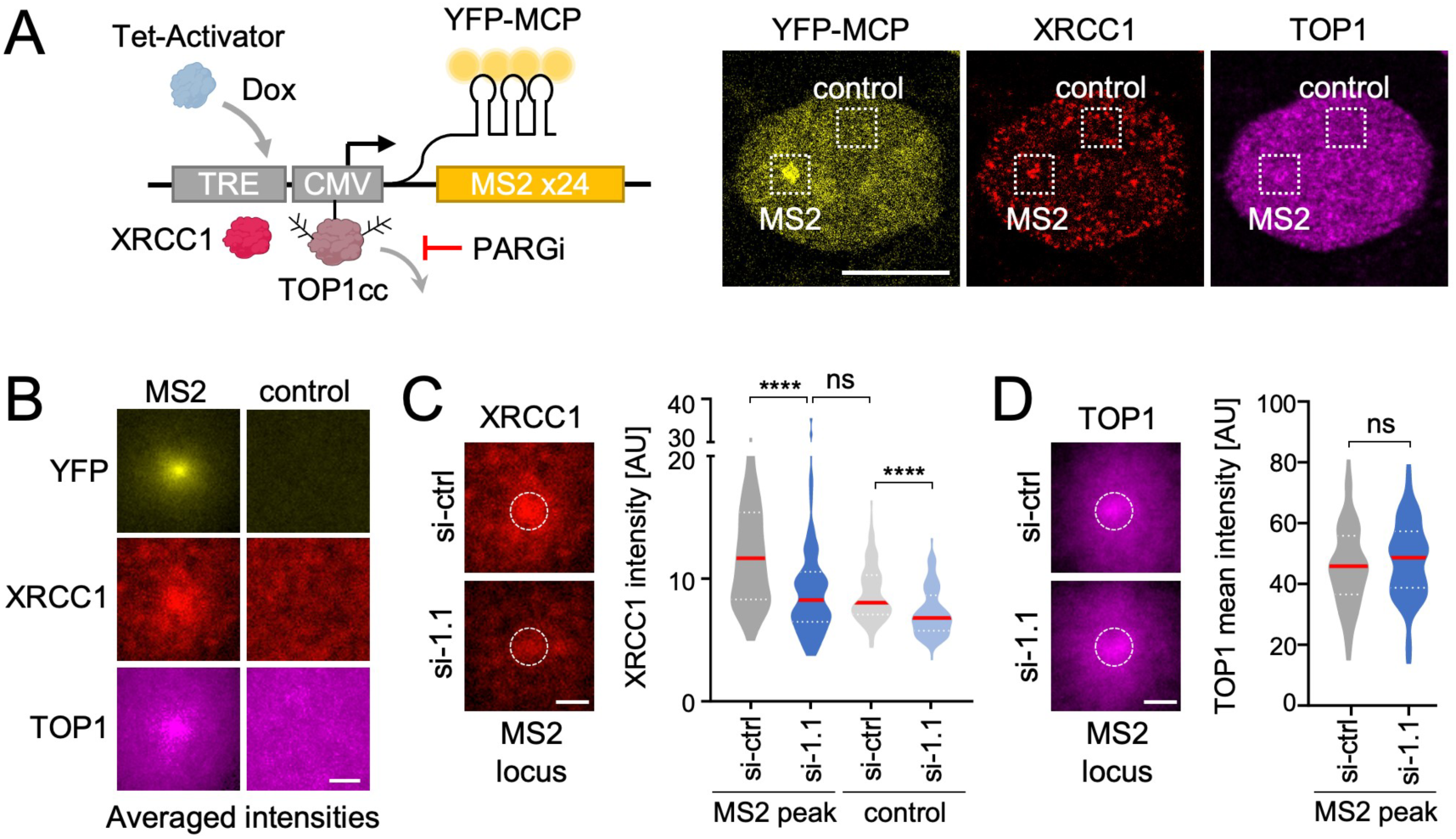
macroH2A1.1 promotes TOP1cc repair at sites of nascent transcription. **(A)** Schematic of U2OS cell-based reporter for transcription-associated DNA damage repair. An MS2 repeat-containing transcript is induced by Dox and detected with YFP-MCP, PARG inhibitor (PARGi) treatment served to stabilize PAR chains. Representative YFP-MCP, XRCC1 and TOP1 IF images of nucleus 5 h after Dox-treatment are shown, PARGi was added for 30 min. Squares depict the MS2 site or a control region used for analyses in (B-D), scale bar: 10 µm. **(B)** Averaged signal intensities (n=82) for the indicated proteins at MS2 or control regions following MS2 induction as in (A). **(C)** XRCC1 intensity distributions 5 h after Dox/PARGi treatment in cells transfected with non-targeting control (si-ctrl), or macroH2A1.1 siRNA (si-1.1). For each MS2 site, mean fluorescence intensity was measured at the MS2 peak (white circle) or a corresponding control region as defined in (A) (n=82). A representative of two independent experiments is shown; si-ctrl cells were used for analyses in (B). **(D)** Average TOP1 intensities and corresponding violin plots as in (C) (n>55). TOP1 signal was measured following 1 h of Dox treatment to avoid potential changes in TOP1 retention due to prolonged repair activity. For all violin plots, red lines reflect the median, P values are based on Mann-Whitney U test; **** p < 0.0001, ns: not significant. Scale bars in B-D are 1 µm.

Next, we assessed the impact of macroH2A1.1 loss on XRCC1 accumulation at MS2 foci. SiRNA-mediated macroH2A1.1 depletion caused a significant reduction in XRCC1 intensity at MS2 peaks without decreasing MS2 mRNA induction or overall XRCC1 protein levels (**Fig. 5C, Fig. S5A-C**). MS2 peak-associated TOP1 accumulation, on the other hand, was comparable in macroH2A1.1-depleted and control cells, supporting the notion that macroH2A1.1 loss does not alter transcription-associated TOP1 engagement but rather the repair events downstream (**Fig. 5D**). Together, these findings demonstrate that macroH2A1.1 regulates XRCC1 function not only upon genotoxic TOP1 inhibition, but also in response to endogenous DNA transactions.

### macroH2A1 splicing is a marker for TOP1i sensitivity in cancer cells

Defects in TOP1cc repair present a potential cancer vulnerability ^38,39^. Exploiting the cytotoxic effects of TOP1cc-associated genome instability, TOP1 inhibition is standard of care for several difficult to treat cancers, including metastatic breast cancer and relapsed ovarian cancer ^40^. However, the efficacy of CPT and its derivatives varies significantly across cancer types and individuals, and the factors that determine TOP1i responsiveness remain largely unknown. Building on the above observations and the fact that macroH2A1.1 splicing is extensively deregulated in cancer ^10,41^, we asked whether variability in macroH2A1.1 expression can predict TOP1i responsiveness. Cancer-related macroH2A1.1 isoform variability is recapitulated in NCI60 breast cancer cells, which display up to six-fold changes in macroH2A1.1 protein levels, while total macroH2A1.2 levels remain largely unchanged (**Fig. 6A**) ^19^. Taking advantage of a large-scale NCI drug-screening program assessing sensitivity of the NCI60 cancer cell line panel to several thousand compounds including > 200 known DNA damaging agents ^42^, we correlated macroH2A1.1 expression with sensitivity to 72 TOP1 inhibitors in the breast cancer subset. Consistent with macroH2A1.1 function as an effector of TOP1cc repair, we observed a robust, inverse correlation between macroH2A1.1 protein levels and TOP1i sensitivity, while TOP1 expression was comparable across cell lines (**Fig. 6A, B**). Sensitivity to two unrelated groups of chemotherapeutics, tubulin inhibitors and HDAC inhibitors, did not significantly correlate with macroH2A1.1 (**Fig. 6B, Fig. S6A**). Using a colorimetric cell viability assay, we confirmed differential sensitivity to CPT treatment in macroH2A1.1^high^ MDA-MB-231 cells and macroH2A1.1^low^ MCF7 cells (**Fig. 6C**). Given that all macroH2A1.1^high^ NCI60 breast cancer cell lines were of the triple-negative subtype (TNBC), we expanded our analysis to include MDA-MB-453 TNBC cells, which express ∼8 times less macroH2A1.1 than their NCI60 TNBC counterparts (**Fig. 6A**). MDA-MB-453 cells were significantly more sensitive to CPT treatment than macroH2A.1.1^high^ MDA-MB-231 cells, supporting the notion that altered TOP1i sensitivity is not merely a consequence of breast cancer subtype (**Fig. 6C**). Providing a mechanistic rationale for increased TOP1i resistance in cancer cells with elevated macroH2A1.1 expression, RADAR analysis revealed faster turnover of CPT-induced TOP1cc lesions in MDA-MB-231 cells compared to macroH2A1.1^low^ MCF7 cells (**Fig. 6D**), and TOP1cc clearance in the latter could be improved by macroH2A1.1 overexpression (**Fig. S3C**). Conversely, macroH2A1.1 depletion increased both TOP1cc levels (**Fig. 2F**) and CPT cytotoxicity in clonogenic and colorimetric survival assays (**Fig. 6E, F, Fig. S6B**). A similar effect was observed in SKOV3 ovarian cancer cells, demonstrating that macroH2A1.1-mediated TOP1cc resistance is not limited to breast cancer cells (**Fig S6C, D**).

**Figure 6.**
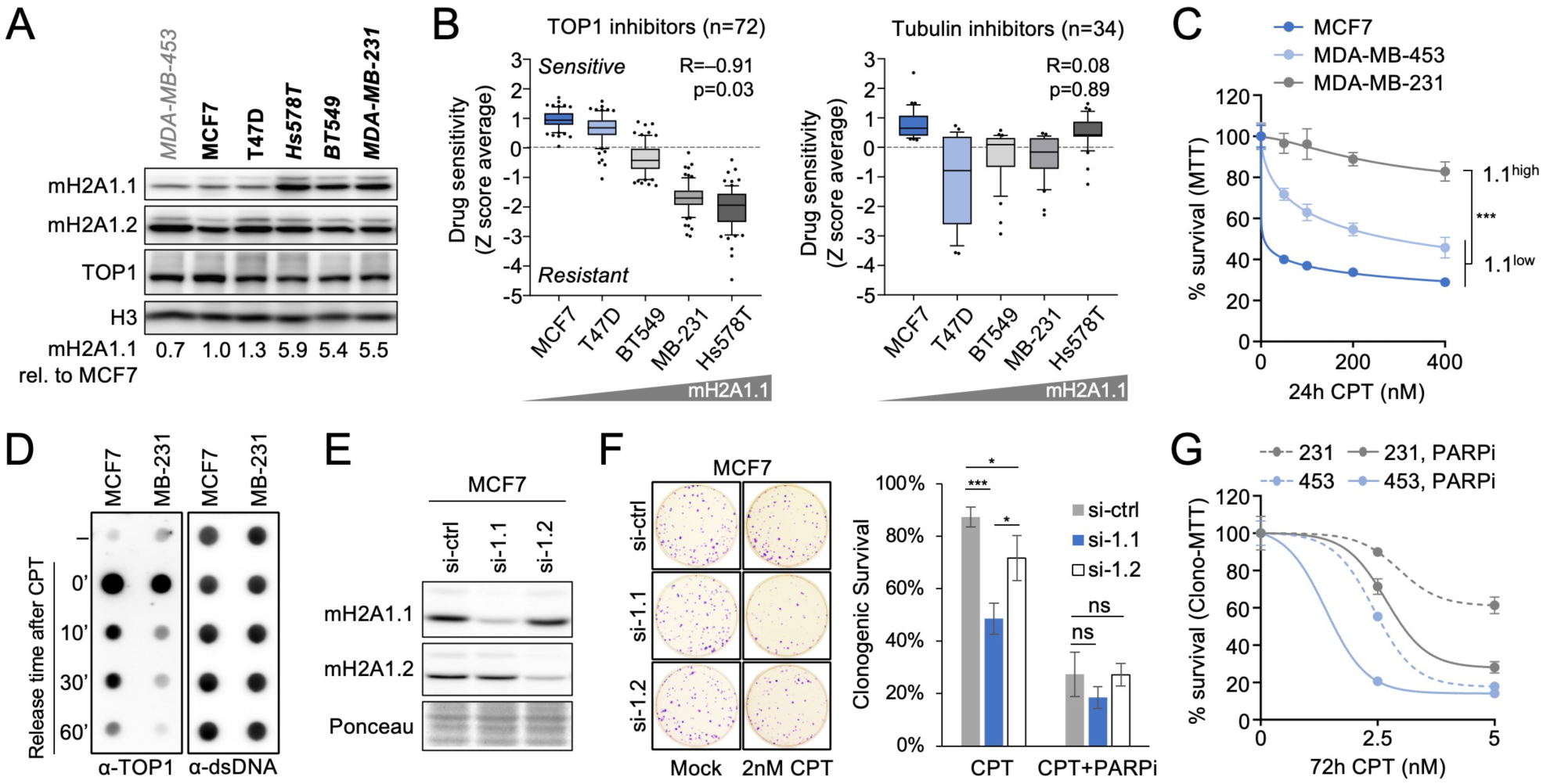
macroH2A1.1 drives TOP1i resistance in cancer cells. **(A)** Western blot for the indicated proteins in MDA-MB-453 and NCI60 breast cancer cell lines (bold). TNBC cell lines are italicized. **(B)** Drug activity levels based on NCI60 drug screen in the indicated breast cancer cell lines, arranged by increasing macroH2A1.1 expression; n=number of compounds tested per drug target. P values are based on Pearson’s Correlation Coefficient. Note that MDA-MB-453 cells were not part of the NCI60 data set. **(C)** Cell viability of the indicated cell lines in response to CPT treatment measured by MTT assay, macroH2A1.1^high^ cells are in gray, macroH2A1.1^low^ cells in blue. Data are presented as mean and SD (n=4); p < 0.001 based on Students two-tailed t-test, relative to MDA-MB-231 cells. **(D)** TOP1 RADAR assay for the indicated cell lines as in Fig. 2F. **(E)** Western blot in MCF7 cells expressing siRNAs against macroH2A1.1 (si-1.1), macroH2A1.2 (si-1.2) or a control sRNA (si-ctrl). **(F)** Clonogenic survival of cells from (E) in response to the indicated drug combinations. Survival was normalized to untreated cells for each siRNA transfection. Representative images are shown, data are presented as mean and SD (n=3). P values are based on Student’s two-tailed t-test, * p < 0.05, *** p < 0.001, ns: not significant. **(G)** Cell viability in the indicated cell lines in response to CPT treatment in the presence or absence of PARPi, measured by MTT after 10 days of clonogenic growth, data are presented as mean and SD (n=3).

Cancer cell sensitivity to TOP1 trapping agents is thought to result primarily from replication stress when DNA polymerase encounters unresolved TOP1ccs. Both replication stress and TOP1cc repair defects can be exacerbated via inhibition or genetic inactivation of PARP1 ^38^. We thus asked whether PARPi treatment can overcome relative CPT resistance in TOP1cc repair-proficient macroH2A1.1^high^ cancer cells. Co-treatment of MDA-MB-231 cells with the PARPi Olaparib resulted in a dose-dependent increase in CPT-induced cytotoxicity that was comparable to the cytotoxicity observed in macroH2A1.1^low^ MDA-MB-453 cells treated with CPT alone (**Fig. 6G**). Of note, macroH2A1.1 depletion did not further aggravate the effect of PARPi in the presence of TOP1i, underlining its PARP/PAR-dependent role in TOP1cc repair (**Fig. 6F**, **Fig. S6D**). Consistent with a macroH2A1.1 isoform-specific TOP1cc repair defect, depletion of the PAR-binding-deficient macroH2A1.2 isoform had only a minor impact on CPT-induced cytotoxicity, which was likely attributable to low levels of replication stress exacerbated by the absence of macroH2A1.2 ^43^ (**Fig. 6F**). Together, these findings point to high macroH2A1.1 expression as a marker for TOP1i resistance, which can be overcome by simultaneously inactivating PARP.

### MacroH2A1.1 expression has predictive value in cancer

To determine potential clinical relevance of our findings, we sought to correlate macroH2A1.1 isoform expression with SSL repair capacity and TOP1i sensitivity in cancer patients. The Single Base Substitution signature SBS2 has been attributed to aberrant BER at sites of cytidine deamination and is frequently observed in breast cancer ^44,45^. We thus compared SBS2 scores derived from whole-genome sequencing data of TCGA breast tumors to macroH2A1.1 splicing efficiency based on matched RNA-Seq data. Consistent with our cell-based findings, we observed a significant inverse correlation between macroH2A1.1 splicing and the SBS2 signature in breast tumor samples with high mutation burden. A similar trend was observed for a second BER-related mutation signature, SBS13 (**Fig. 7A**). To evaluate implications for TOP1i-based cancer therapy, we asked whether macroH2A1.1 expression, and by extension SSL repair capacity, can affect survival outcomes in patient groups treated with TOP1 trapping agents. Using the Kaplan-Meier Plotter survival analysis tool ^46^, we identified a subset of TCGA ovarian cancer patients with macroH2A1.1 splice variant-specific gene expression information that was treated with the TOP1i topotecan ^47^. Low macroH2A1.1 expression was significantly correlated with improved survival outcome in this patient group. Conversely, macroH2A1.1 levels were not indicative of survival outcome in ovarian cancer patients treated with the microtubule stabilizer taxol (**Fig. 7B**), consistent with our cancer cell line data (**Fig. 6B**), or the nucleoside analog gemcitabine (**Fig. S6E**). Altogether, we find that macroH2A1 splicing state can be indicative of SSL repair proficiency in cancer, which may translate into a protective role for macroH2A1.1 in response to TOP1i treatment and adversely affect treatment responsiveness.

**Figure 7.**
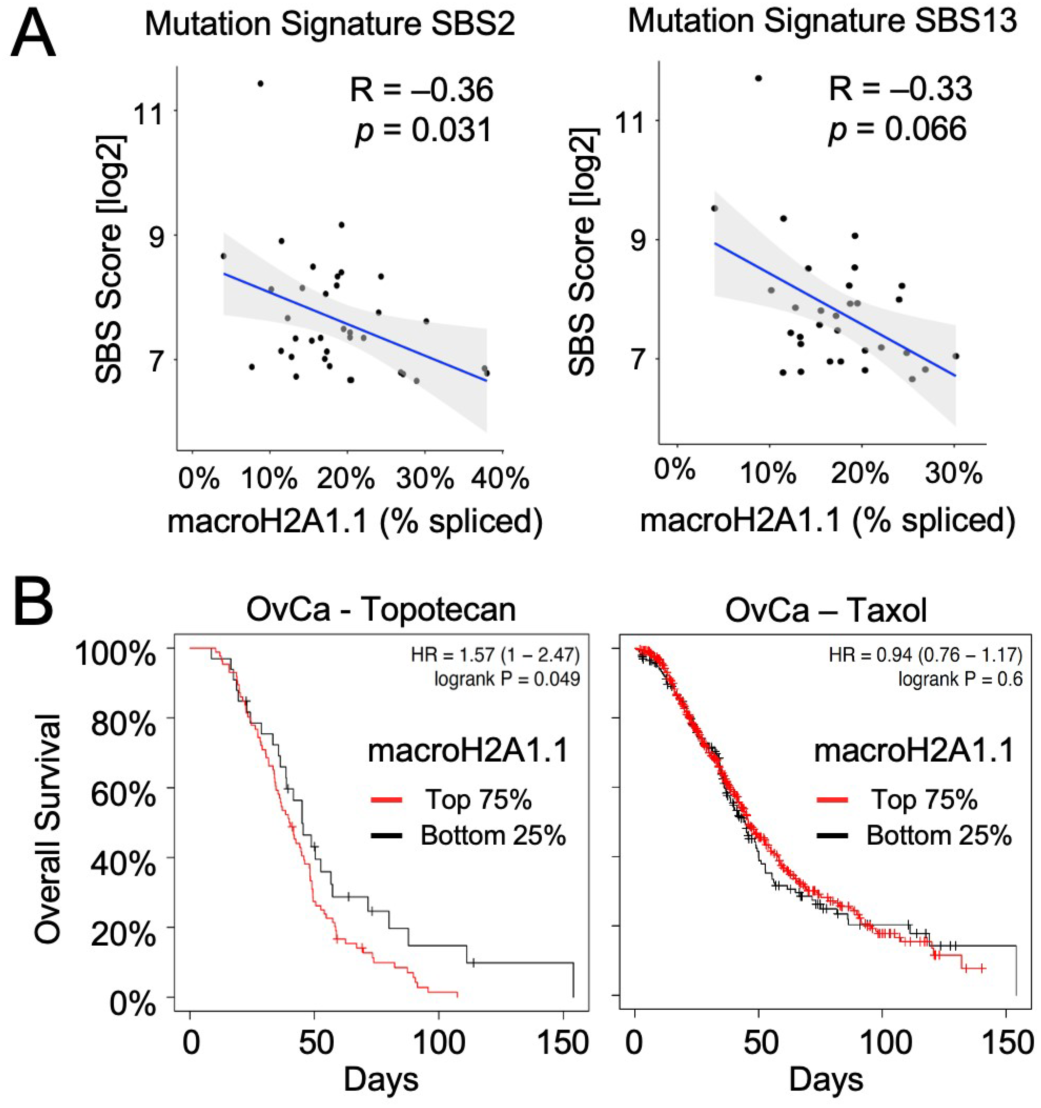
macroH2A1.1 expression is indicative of SSL resistance in cancer patients. **(A)** Correlation between relative macroH2A1.1 expression and BER-related APOBEC signature scores (SBS2 and SBS13) in TCGA breast tumors with high mutation burden (SBS scores > 100). P values are based on Spearman correlation. **(B)** Kaplan-Meyer survival analysis of TCGA ovarian cancer patient subgroups where treatment regimens contained Topotecan (n=119) or taxol (n=821). Patients were stratified by macroH2A1.1 mRNA expression based on an isoform-specific Affymetrix microarray probe (214500_at), the bottom 25% were considered macroH2A1.1 low expressors.

## Discussion

We have identified macroH2A1.1 as a chromatin effector of TOP1-associated genome maintenance. By coordinating PAR/PARP-dependent repair processes, macroH2A1.1 facilitates the resolution of torsional constraints while simultaneously protecting from aberrant TOP1cc accumulation. This function has clinical implications for therapeutic regimen involving TOP1 inhibition and points to macroH2A1.1 as a predictive marker for cancer cell sensitivity to TOP1 poisons.

Our findings place macroH2A1.1 at the nexus of the TOP1-PARP1 axis that regulates the cellular response to topological stress. Reminiscent of the enrichment of the HR-promoting macroH2A1.2 variant at sites of recurrent replication stress ^43^, we found that macroH2A1.1 colocalizes with TOP1 domains across the genome. While chromatin remodeling factors have previously been implicated in enhancing TOP1 accessibility to DNA ^48^, macroH2A1.1 adds a level of epigenetic control that is directly linked to TOP1 activity, concomitant PARP activation and base excision repair factor recruitment. Consistent with this, macroH2A1.1 not only co-occupies TOP1-enriched chromatin domains but interacts with TOP1 in a DNA damage and PARP-dependent manner. By mediating TOP1cc repair at hotspots of TOP1 function, macroH2A1.1 thus provides a rationale for the seeming discrepancy between TOP1 enrichment and concomitant TOP1cc depletion at sites of recurrent topological stress ^17,26^. Although it is conceivable that PARP/PAR helps direct the targeted chromatin incorporation of macroH2A1.1 at TOP1 peaks, we find that macroH2A1.1 accumulation does not require its PAR binding domain (**Fig. S3B**). What controls macroH2A1.1 deposition at sites of TOP1 activity will be an important subject of future investigation. Taken together, our observations provide a rationale for promoter-associated macroH2A1.1 enrichment beyond gene regulation that involves the maintenance of genome integrity. They further suggest that gene expression changes observed upon macroH2A1.1 loss may at least in part be attributed to impaired resolution of torsional stress ^14,15^.

Given that macroH2A1.1 interacts with PARP1 in a PAR-dependent manner, we propose that auto-PARylated PARP1 may serve as a bridge between macroH2A1.1 nucleosomes and BER factors with conserved PAR binding motifs such as XRCC1 ^31^. This function is likely further enhanced through macroH2A1.1-mediated PAR chain stabilization and concomitant changes in the underlying chromatin structure ^20,49^, and is consistent with the reciprocal relationship between PARP1 and XRCC1, wherein recruitment of one directly depends on the other ^32,49^. Using a cell-based reporter for the detection of nascent transcripts, we demonstrate that macroH2A1.1-mediated control of XRCC1 recruitment is relevant in the context of transcription-induced topological stress, which may at least in part explain the TSS-associated increase in TOP1cc levels observed when macroH2A1.1 is depleted (**Fig. 2C**, **Fig. 5C, D**). We anticipate that imaging-based interrogation of TOP1-associated repair factors, including but not limited to XRCC1, will significantly advance our understanding of the molecular events that underlie the highly dynamic resolution of torsional constraints. Together with its recently described roles in MMEJ and oxidative lesion repair ^13,49^, our findings firmly establish macroH2A1.1 as a chromatin effector of PAR-dependent DNA repair.

Given the distinct repair functions of macroH2A1 splice isoforms ^13,43,49–51^, we propose alternative macroH2A1 splicing as a means to adjust the chromatin environment to changing genome maintenance needs. MacroH2A1.1 expression was found to increase upon cellular differentiation and in in postmitotic cells ^10,12,52^, where the accumulation of SSLs is a main source of deleterious damage that is often associated with degenerative diseases ^1,53^.

Conversely, expression of the HR-promoting, replication stress-protective macroH2A1.2 isoform is elevated in replicating cells, including many cancers ^10,12^. The two isoforms may furthermore play complementary roles in solving the problem of topological stress, as TOP1cc repair can involve homologous recombination during S phase, owing to the formation of DSBs that are produced by the collision of replication forks with unresolved TOP1ccs ^3^. Consistent with this notion, both macroH2A1 isoforms often co-occupy the same chromatin domains ^14–16,43^, and two chaperones recently identified to promote macroH2A1 chromatin deposition do not distinguish between macroH2A1 splice isoforms ^54,55^. It will nevertheless be interesting to investigate if macroH2A1.1-specific control of TOP1cc repair is linked to a distinct and/or unique chromatin remodeling pathway.

Aberrant TOP1cc accumulation has been linked to various genome instability syndromes and is exploited as a genotoxic vulnerability in cancer therapy ^2,5,6^. Our findings in cancer cell lines and TCGA patient samples suggest that macroH2A1.1 expression is predictive of TOP1cc repair capacity, which may ultimately help refine TOP1i treatment strategies. For example, macroH2A1.1^high^ cancers may benefit from TOP1i/PARPi combination treatment, whereas macroH2A1.1^low^ cancers appeared inherently susceptible to TOP1i alone (**Fig. 6G**). It will further be interesting to determine if manipulation of macroH2A1.1 splicing can help enhance TOP1i sensitivity. Beyond cancer treatment, macroH2A1.1 may protect from repeated TOP1 cleavage, particularly at actively transcribed genes, which were recently shown to accumulate a unique TOP1 mutation signature that contributes to mutagenesis and malignant transformation ^5,56^.

Altogether, we have uncovered macoH2A1.1 as an epigenetic rheostat of TOP1 activity that ensures genome integrity by establishing a TOP1-permissive chromatin environment. Deregulation of this pathway is tied to sensitivity to TOP1 DNA lesions and may be therapeutically exploited as a cancer vulnerability.

## Data Availability

All genomic data will be made available for health/medical/biomedical research use via the database of Genotypes and Phenotypes (dbGaP) under accession number phs003729.

## Acknowledgements

We thank Yves Pommier for critical reading of this manuscript, Yan Zhang and Ludmila Danilova (JHU Experimental Cancer Genomics Core) for help with mutation signature analyses, Michael Matunis and Allison DeHaas (JHU Bloomberg School for Public Health) for microscope assistance, Anne-Claire Lavigne and Kerstin Bystricky (University of Toulouse) for MDA-MB231 RNA-Seq data, and Roger Greenberg (University of Pennsylvania), Daniele Gilkes (JHU) and Tian-Li Wang (JHU) for reagents. This work was supported by internal Johns Hopkins University funding and the National Institutes of Health National under award numbers R35GM153484 and R01CA285725 (PO and THL), by Knut och Alice Wallenbergs Stiftelse: KAW 2016.0161, KAW 2022.0380 and KAW 2022.0189, Vetenskapsrådet: 2021-02630 VR, Cancerfonden: 21 1771 Pj01 H, and Karolinska Institutet Consolidator: 2-190/2022 (LH and VK), and the national grant PID2021-126907NB-I00 from MCIN/AEI/10.13039/501100011033, co-funded by European Regional Development Fund (MB and DC).

## Author Contributions

PO, THL and CXQ designed the experiments. THL performed all cell-based and biochemistry assays. CXQ and TW performed sequencing experiments, CXQ and VK analyzed genomics data. YS performed MS2 reporter cell assays, VR and XZ generated CRISPR knockout cells. DC and MB provided macroH2A1 expression vectors. PO and THL wrote the manuscript with input from LB, MB and CXQ.

## Supplemental information

Figures S1–S6, appended

Document S1. Table S1

## Experimental Model and Subject Details

### Cell Lines

Human breast cancer cell lines MCF7 (ATCC), MDA-MB-231 (ATCC), MDA-MB-453, Hs-578T, T47D and BT-549 (gift from D. Gilkes, JHU), as well as HEK293 T-Rex, HEK293 cells (gift from The Broad Institute, Cambridge) and U2OS 2-6-3 cells (gift from R. Greenberg, U. Penn) were grown in Dulbecco’s modified eagle medium (DMEM, Invitrogen) supplemented with 10% FBS (Gemini) and maintained at 37 °C with 5% CO_2_. All cell lines were regularly tested for mycoplasma using Mycoplasma PCR detection kit (Abcam). For transient knockdown, siRNAs were transfected using DF-1 reagent following the manufacturer’s instructions (Dharmacon) and analyzed at 72 - 96 h post transfection. For stable knockdown, lentiviral infection of LKO.1 shRNA-expression vectors was carried out by spin infection (2500 rpm, 90 min, Eppendorf 5810R centrifuge) with 8 µg/ml polybrene (Sigma). Cells were incubated overnight prior to virus removal and selection with puromycin (1-2 µg/ml, Invitrogen). CRISPR/Cas9 knockout of macroH2A1.1 exon 6b was preformed using two exon-flanking guide RNAs. In brief, crRNA, tracrRNA and GFP-tagged SpCas9 protein were assembled into an RNP complex, followed by nucleofection (Amaxa Nucleofector) and single cell sorting of GFP^+^ cells. Clones were screened by PCR for homozygous gene inactivation, cells with intact exon 6b alleles served as clonal wildtype controls. See Table S1 for sequence information.

### Plasmids

pLKO.1-puro-based shRNA expression vectors were published previously or generated following the provider’s instructions (Addgene), see Table S1 for shRNA sequences. pLVX-FLAG-macroH2A1.1 vector was published previously ^20^, pLVX-FLAG-macroH2A1.1 G224E was generated from the latter by replacing a 1176 bp XhoI/XbaI fragment with a G224E (GAG>GGT) mutation-containing gBlock using Gibson cloning.

## Method Details

### Rapid approach to DNA adduct recovery (RADAR)

RADAR was performed essentially as described previously ^28^. 1 x 10^6^ cells were treated with 1 µM CPT for 30 min, and then released into media for the indicated times. Cells were then washed with 1 x PBS (GIBCO) and lysed with 600 µL DNAzol (Invitrogen), followed by ethanol precipitation. DNA pellets were solubilized in 8 mM NaOH at 4 °C overnight, then heated at 65 °C for 5 min, followed by shearing with sonication (Fisher Scientific Sonic Dismembrator). DNA content was quantified using NanoDrop One (ThermoFisher), diluted with 25 mM NaPO_4_ (pH 6.5) to 1-2 μg of DNA per sample, then vacuum-blotted onto nitrocellulose membrane (GE Biosciences) using the Bio-Dot apparatus (Bio-rad). The membrane was washed with 2x SSC, dried and irradiated with 120 mJ/cm^2^ using a UV Stratalinker 2400 (Stratagene). The membrane was immunostained using TOP1 and dsDNA antibodies. HRP-conjugated secondary antibodies were used for signal detection by enhanced chemiluminescence (Advansta). Images were captured with the ChemiDoc MP imaging system (Bio-Rad).

### Alkaline comet assay

The alkaline comet assay was performed according to the Trevigen CometAssay™ kit protocol. Cells were treated with 20 μM CPT for 1 h, then trypsinized at 37 °C for 2 min. The cells were centrifuged and resuspended in ice-cold PBS and 500,000 cells/mL were mixed with LMAgarose (R&D systems) at 37 °C at a ratio of 1:10 (v/v). The cell/agarose mixture was transferred onto CometSlides (R&D systems), solidified and immersed in prechilled lysis solution (R&D systems) overnight at 4 °C. For DNA denaturation, the slides were immersed in alkaline unwinding solution (200 mM NaOH, 1 mM EDTA) for 20 min at RT, followed by electrophoresis in alkaline electrophoresis solution (200 mM NaOH, 1 mM EDTA) at 4 °C. After electrophoresis, the slides were washed twice in dH_2_O, once in 70 % EtOH, dried and incubated with SYBR® Gold in TE buffer for 30 min. The slides were mounted with ProLong Gold Antifade Mountant (ThermoFisher) and imaged using EVOS M7000 Imaging System (Invitrogen). Statistical analysis was performed using OpenComet in ImageJ.

### Immunofluorescence imaging and analysis

For immunofluorescence (IF), MCF7 or U2OS 2-6-3 cells were plated on poly-L-lysine-coated coverslips 24 h prior to the respective treatments and fixed with 4% formaldehyde for 10 min at RT. Cells were permeabilized with 0.5% Triton-X in 1x PBS for 10 min, washed with 1x PBS-T (0.1% Triton-X) and blocked with 20% FBS in 1x PBS-T for 1 h at RT. After blocking, cells were washed and incubated with primary antibodies against XRCC1 (Mouse anti-XRCC1, Santa Cruz sc-56254, diluted at 1:300) and/or TOP1 (Rabbit anti-TOP1, Abcam ab109374, diluted at 1:300) for 1 h at 37 °C. Cells were washed and incubated with secondary antibodies diluted in 1x PBS-T, 5% FBS for 1 h at RT. Coverslips were immersed in 1x PBS with 5 µg/mL of Hoechst 33342 (Sigma-Aldrich) for 5 min and mounted on slides with ProLong Gold Antifade Mountant (ThermoFisher). For S phase staining, MCF cells were pulse-labeled with 10 µM EdU for 30 min. Where indicated, cells were subsequently treated with 1 µM CPT treatment for 30 min. After fixation and permeabilization, cells were subjected to Click chemistry labeling (Click-It Kit, Invitrogen) and stained with 5 μM AZDye 647 Azide Plus (Vector Laboratories) for 30 min at RT. Cells were then stained with XRCC1 antibody as described above. Images were captured with Zeiss microscopes (Axio Imager Z1 or LSM 900 with Airyscan2) and analyzed using ImageJ. For XRCC1 foci analysis, at least 300 nuclei were randomly collected and nuclear foci were identified using the ‘Find maxima’ function in ImageJ. For MS2 intensity analyses, a ∼4×4 µm square was centered on the MS2 peak or a randomly selected MS2-distal nuclear control region, at least 50 regions were collected from each group following MS2 induction with doxycycline (2 µg/mL, 1 h or 5 h). Cells were treated with 10 µM PARGi PDD 00017273 (MedChemExpress) for 30 min prior to sample collection to prevent PAR chain degradation. “Image sequence” function (ImageJ) was used to combine images from the same color channel into a single TIFF file. To determine TOP1, XRCC1 or MS2 signal intensities at the MS2 peak, mean intensities within a 1 µm-wide circle centered on the MS2 peak were measured for all stacked images. Corresponding nuclear background was defined as the average intensity within the 4×4 µm control region of the same cell.

### Cellular extract preparation and immunoblotting

10^6^ cells were lysed in RIPA lysis buffer (25 mM Tris-HCl, pH 7.5; 150 mM NaCl; 2 mM EDTA; 1% NP-40; 1% Na-deoxycholate; 0.1% SDS) supplemented with cOmplete^TM^ EDTA-free protease inhibitor cocktail (Roche). Lysates were sonicated, centrifuged, diluted with 5x sample buffer (312.5 mM Tris-HCl, pH 6.8; 10% SDS; 50% glycerol; 12.5% β-mercaptoethanol; 0.05% bromophenol blue) and heated for 10 min at 95 °C. Lysates of equal protein amount based on BSA assay (Bio-Rad) were separated by SDS-PAGE and subjected to western blotting using the indicated primary antibodies. HRP-conjugated secondary antibodies were used for signal detection by enhanced chemiluminescence (Advansta). Images were captured with the ChemiDoc MP imaging system (Bio-Rad).

### Co-immunoprecipitation

HEK293 T-REx cells expressing Flag-tagged macroH2A1.1 or macroH2A1.2 used for co-IP experiments were described previously ^13^, parental cells were used as a control. 10^7^ cells were treated as outlined in the figure legends, CPT treatment was at 1 µM for 30 min, PARPi at 10 μM for 24 h. For co-treatment of CPT and PARPi, cells were pretreated with 10 μM PARPi for 30 min, followed by CPT treatment. Equal numbers of cells were used for each IP experiment. Briefly, cells were washed with PBS and resuspended in hypotonic buffer (20 mM HEPES; 10 mM KCl; 2mM MgCl_2_; 10% glucose; 0.5% NP-40) followed by MNase digestion (0.2 U/μl, 4°C, 60 min) in digestion buffer (20 mM HEPES; 150 mM KCl; 10% Glycerol; 3 mM CaCl_2_). 20 mM EGTA was used to terminate the reaction. Samples were homogenized using a 27G needle, followed by centrifugation at 1000 g, retrieval of the supernatant and IP with M2 magnetic beads (50% slurry) at 4°C for 60 min. Post IP, beads were washed 4 times using wash buffer (20 mM HEPES; 150 mM KCl; 0.1% NP-40) followed by 50 mM HEPES. Immunoprecipitated proteins were eluted by incubating beads with SDS sample buffer containing 10% β-mercaptoethanol for 5 min at 95°C, followed by western blotting.

### Cell viability assays

For clonogenic survival assays, cells were seeded at 300 cells per well in 6-well Evap EDGE plates 8-16 h prior to treatment with the indicated doses of CPT with or without 0.1 µM Olaparib. After 72 h treatment, cells were released from drugs, maintained for 7-10 days, and then fixed with staining solution (6% glutaraldehyde and 0.5% crystal violet). Colonies were counted by Scan 4000 (Interscience). For MTT assays, cells were seeded at 2,000-4000 cells per well in 96-well plates and treated with the indicated doses of CPT for 24 h. 48-72 h after the end of drug treatment, cells were treated with 0.5 mg/mL MTT reagent (Sigma Aldrich) for 2 h, washed and incubated with 100 µL DMSO for at least 20 min. Absorbance was measured at 570 nm using SpectraMax M5 (Molecular Devices), with SoftMax Pro 5.2 software (Molecular Devices). A modified MTT assay was performed to assess long-term cell survival in cell lines with impaired colony formation. 10 days after the end of drug treatment, cells were stained with 0.5 mg/mL MTT for 2 h, washed and incubated with 1-2 mL DMSO for at least 20 min. 100 µL of supernatant from each sample was transferred to 96-well plates and processed as above.

### RNA Extraction and RT-PCR

Total RNA was extracted using the TRIzol^TM^ reagent according to the manufacturer’s instructions (Invitrogen). cDNA was synthesized from 2 µg of total RNA using PrimeScript RT Master Mix (Takara), and expression of the indicated genes was analyzed by quantitative RT-PCR using the CFX Opus 96 Real-Time PCR System (Bio-Rad) (see Table S1 for primer sequences).

### Cleavage Under Targets & Release Using Nuclease (CUT&RUN)

CUT&RUN was performed essentially as described with minor modifications ^57^. Concanavalin A-coated (ConA-coated) bead slurry (Bangs Laboratories, cat. BP531) was activated in binding buffer (20 mM HEPES, pH 7.5; 10 mM KCl; 10 mM CaCl_2_; 10 mM MnCl_2_). 10^6^ cells were harvested and resuspended in 3 ml wash buffer (20 mM HEPES pH 7.5; 150 mM NaCl; 0.5 mM Spermidine; Complete Protease inhibitor (Sigma-Aldrich)) at RT. 10 µL of activated bead slurry was added and the samples were rotated for 10 minutes at room temperature. Bead-bound cells were isolated on a magnet stand and resuspended in 300 µL antibody buffer (dig-Wash buffer: wash buffer plus 0.05% digitonin, supplemented with 2 mM EDTA). 0.3 µg of each primary antibody was added per sample and incubated under rotation at 4 °C overnight. Beads were washed and resuspended in 300 µL dig-Wash buffer. 5 µL pAG-MNase (EpiCypher) was added and bead slurries were rotated at 4 °C for 1 h. Samples were washed twice in dig-Wash buffer and MNase was activated via addition of 2 mM CaCl_2_, followed by a 1 h incubation at 0 °C. MNase was inactivated with 2x STOP Buffer (340 mM NaCl; 20 mM EDTA; 4 mM EGTA; 0.05% digitonin; 100 µg/mL RNAse A; 50 µg/mL glycogen) mixed with 1-2 ng CUTANA E. coli Spike-in DNA (EpiCypher). CUT&RUN fragments were released via incubation at 37 °C for 30 minutes, separated on a magnet stand, and purified using the MiniElute PCR Purification Kit (Qiagen). Selected CUT&RUN samples were analyzed by qPCR enrichment using a CFX Opus 96 Real-time PCR System (Bio-rad), see Table S1 for primer sequences.

### TOP1 Covalent Adduct Detection coupled to NGS (TOP1 CAD-Seq)

For the detection of steady-state TOP1ccs, 10^7^ cells were pre-treated with MG132 (10 μM, Sigma-Aldrich) for 30 min to inhibit proteasomal degradation of TOP1ccs, followed by a brief 3-5 min pulse of CPT (20 µM). For detection of TOP1ccs after prolonged damage and repair, cells were treated with 20 µM CPT for 30 min in the absence of MG132. DNA isolation and TOP1 IP was performed essentially as described by Kuzin *et al*. ^25^. Following treatments, cells were immediately lysed and briefly sonicated using a Fisher Scientific Sonic Dismembrator Model 100. DNA covalent adducts were ethanol-precipitated and resuspended in TE-0.1% SDS plus 0.5 mM AEBSF (Sigma-Aldrich). Samples were further sonicated with Covaris ME220 sonicator for 5 min using the High Cell protocol in 1 ml milliTUBEs to produce ∼1 kb-sized fragments. For immunoprecipitation, 2 μg of anti-TOP1 antibody (Abcam) was preincubated with 25 μl of Protein A/G magnetic beads (Pierce) in RIPA buffer at 4°C for 3 hours. DNA covalent adducts from 10^7^ cells was added to the Protein A/G–antibody complexes and incubated overnight at 4°C. Magnetic beads were washed as described ^25^, DNA was eluted in 100 µL of TE-0.5% SDS buffer with Proteinase K at 60 °C for 4 hours and purified using the MiniElute PCR purification kit (Qiagen).

### Library Preparation and Next Generation Sequencing

NGS libraries were prepared using the ThruPLEX DNA-Seq Kit (Takara Bio USA, cat. R400675), following the manufacturer’s instructions. 1-10 ng DNA was used per sample and PCR cycle numbers were adjusted as recommended. Samples were bar-coded using the DNA HT Dual Index Kit (Takara Bio USA). Amplified libraries were purified using AMPure XP beads (Beckman Coulter) at a 1:1 (v/v) ratio. Library fragment size and concentration were measured using TapeStation (Agilent D1000 reagent). Up to 18 samples were pooled at a final concentration of 4 nM per sample. Paired-end Illumina sequencing was performed on the NextSeq 2000 Sequencing System.

### NGS Data analysis

The quality of the raw sequenced reads was examined with FastQC and adapters trimmed using cutadapt. PE reads were aligned against the hg38 human reference genome using Bowtie2 ^58^. Aligned reads were sorted, indexed, deduplicated using Picard (Broad Institute, GitHub) and Samtools ^59^. For CUT&RUN, reads were further aligned against the E-coli (K12_DH10B) genome and human aligned reads were normalized to CUTANA E. coli spike-in content following EpiCypher’s instructions. For CAD-Seq, replicates were merged using Samtools to increase read depth. Fastq and bam files will be made available through dbGaP (phs003729). H3K27me3 (GSM949581) and PARP1 ChIP-Seq data (GSM1517306) were obtained from Gene Expression Omnibus (GEO) ^21,23^. Reads mapping to ENCODE blacklist regions were excluded from downstream analyses ^60^.

#### Peak-calling

FLAG-macroH2A1.1 broad peaks were called using SICER ^61^ with 200 bp window size, 150 fragment size, and 600 bp gap size. TOP1 narrow peaks were called using MACS2 ^62^. The R package ChIP-seeker was used to annotate peaks based on genomic context ^63^. TSS-proximal TOP1 peaks were defined as all TOP1 peaks within +/-3000 bp of the nearest TSS. TSS-distal (Non-TSS) TOP1 peaks were defined as all TOP1 peaks greater than 10 kb away from the nearest TSS, strand orientation was considered only for TSS-proximal peaks.

#### Z-Normalization of BigWig Files

To standardize the genomic signal across different datasets, we performed z-normalization on BigWig files, where indicated. Each score in the BigWig file was transformed into a z-score by subtracting the mean and dividing by the standard deviation: Z-score = (score–mean)/SD.

#### Correlation Analysis

Similarity between CUT&RUN data sets was evaluated using the multiBigwigSummary and plotCorrelation features in deepTools ^64^. The multiBigwigSummary tool was employed to compute average scores across individual bigwig files for different experimental replicates based on equally sized 10 kb bins, plotCorrelation was used to compute and visualize the Spearman correlation coefficients between these datasets.

#### Jaccard Index Analysis

The Jaccard Index provides a measure of similarity between peak sets that is comparable across sets of varying size. Jaccard Indices were calculated using bedtools ^65^, and are defined as the bases in the intersection divided by the bases in the union, producing values bounded by zero and 1 (no overlap and complete overlap, respectively). To determine whether this metric was different than expected by chance, permuted Jaccard values were calculated based on random shuffles of both peak sets (n = 1,000 shuffles) using the shuffle command from bedtools within the bounds of the human genome assembly hg38. All peak sets were called with SICER ^61^.

#### Profile plots and heatmaps

The computeMatrix command from deepTools was used to calculate scores per genome regions defined in a BED file. Heatmaps were generated using the plotHeatmap command, profile plots using the plotProfile command. TSS separation based on gene expression quartiles was based on RNA-Seq data from ^15^.

#### LOESS Smoothing of Average Profiles

To smooth the average profiles of our genomic data, we applied LOESS (Locally Estimated Scatterplot Smoothing) using the geom_smooth function in R’s ggplot2 package. We utilized span = 0.15 to capture localized trends in the data while avoiding overfitting, a span = 0.025 was used for Fig. 2B to provide finer detail.

### Mutation signature and Kaplan-Meyer analyses

SBS2/SBS13 mutation signature analyses were based on Catalog of Somatic Mutations in Cancer (COSMIC) scores derived from the TCGA breast cancer patient data set ^66^. MacroH2A1.1 splicing efficiency was calculated based on matched RNA-Seq data as FPKM macroH2A1.1 (uc003lan.1) divided by the sum of FPKM macroH2A1.1 and FPKM macroH2A1.2 (uc003lam.1, uc003lao.1, uc003lat.1, uc003las.1), isoform IDs are based on UCSC genome browser annotations. Correlation analyses between macroH2A1.1 expression and overall patient survival were performed using the Ovarian Cancer data set in the Kaplan-Meier Plotter survival analysis tool ^46,47^ and the macroH2A1.1-specific Affy probe set 214500_at ^19^. Patient subsets and cutoffs are as indicated in the figure legend, array quality control was set to “exclude biased arrays”.

## Supplemental Figures

**Figure S1, related to Figure 1.**
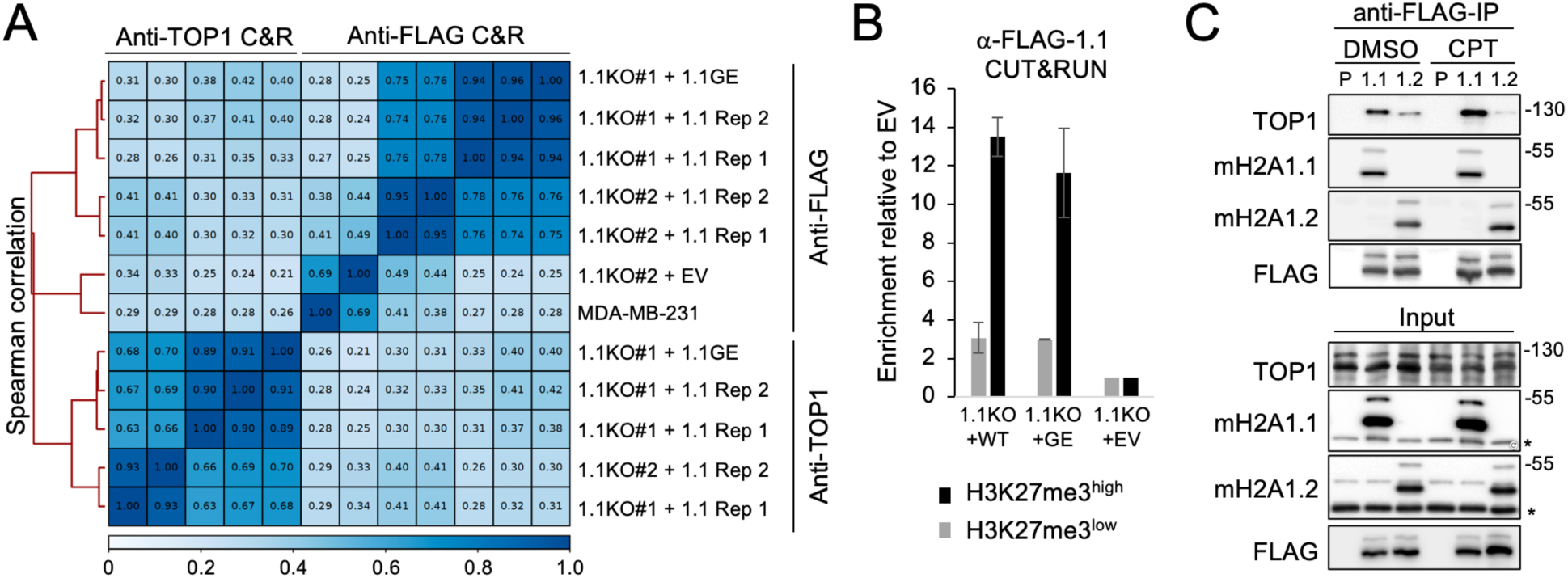
(**A**) Unsupervised hierarchical clustering of the indicated CUT&RUN NGS samples based on Spearman Correlation Coefficients. 1.1KO#1 and 1.1KO#2 represent independent CRISPR knockout clones, Rep 1 and Rep 2 independent experimental replicates. (**B**) qPCR analysis of FLAG-macroH2A1.1 CUT&RUN samples from MDA-MB-231 macroH2A1.1 knockout (1.1KO) cells reconstituted with WT or G224E (GE) mutant macroH2A1.1, or empty vector (EV) at an H3K27me3high heterochromatin region known to bind macroH2A1.1 and an H3K27me3low control region 15, see Table S1 for primer sequences. Samples were normalized to the EV control, values are expressed as mean and SD (n=2). (**C**) Western blot for the indicated proteins in nuclear lysates (input) or anti-FLAG IP samples from parental (P) 293 cells and FLAG-macroH2A1.1 (1.1) or FLAG-macroH2A1.2 (1.2) knock-in cells in the presence or absence of CPT treatment; * endogenous macroH2A1.1 or macroH2A1.2 protein.

**Figure S2, related to Figure 2.**
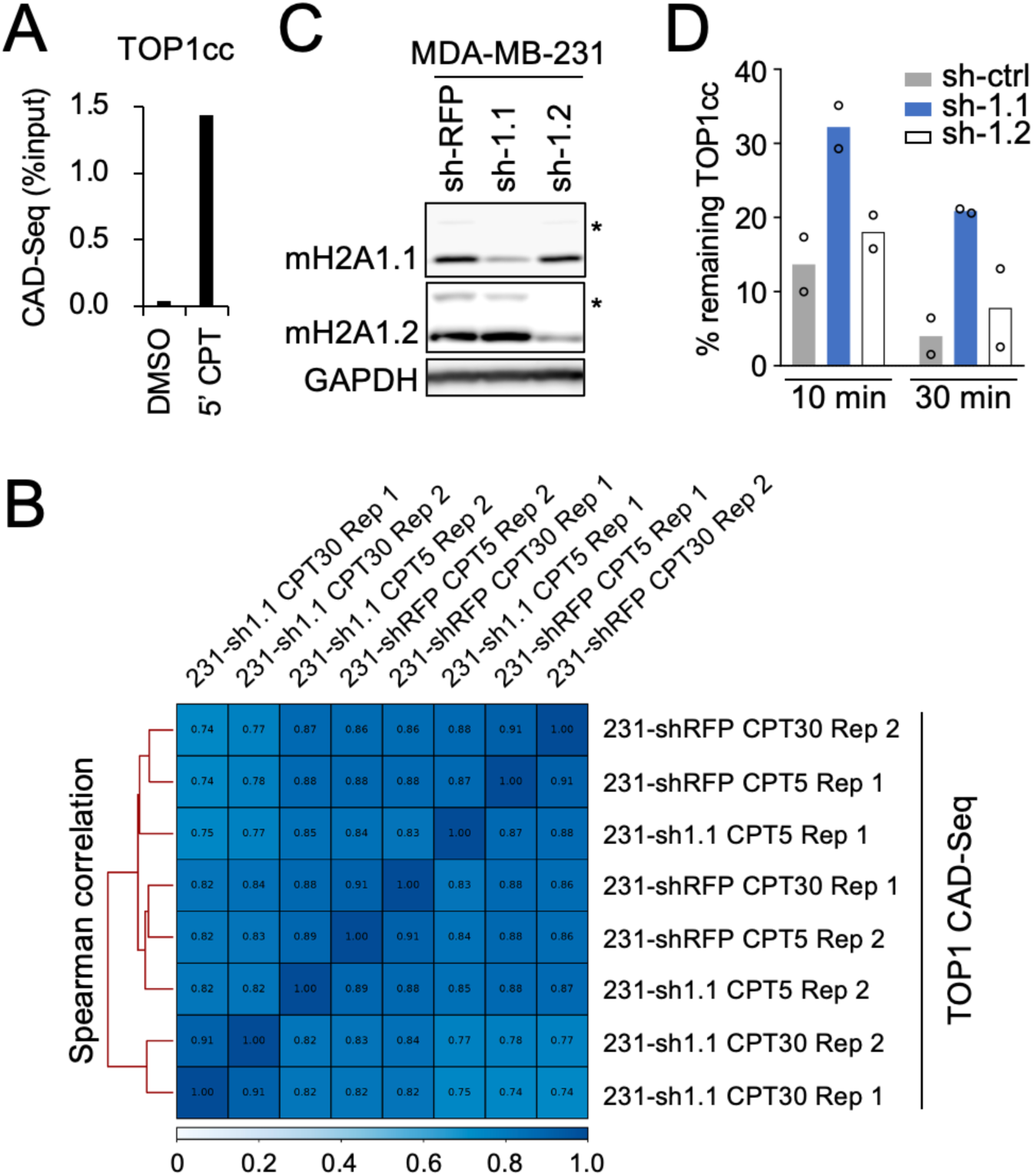
**(A)** qPCR analysis of CAD-Seq IP DNA relative to input at the TOP1cc-rich MYC promoter as described in Ref. 25, in MDA-MB-231 cells in the absence (DMSO) or presence of TOP1cc stabilization (CPT 5’). **(B)** Unsupervised hierarchical clustering of the indicated TOP1 CAD-Seq samples as in Fig S1A. CPT5 and CPT30: 5 min and 30 min CPT treatment, respectively; Rep: replicate experiment. **(C)** Western blot for the indicated proteins in MDA-MB-231 cells stably expressing shRNAs against macroH2A1.1, macroH2A1.2 or RFP (non-targeting control); * ubiquitinated macroH2A1 isoforms. **(D)** Quantification of TOP1cc turnover at the indicated time points after CPT treatment. The percent of remaining TOP1cc relative to 0’ after CPT is shown for two independent RADAR assays (open circles), see Fig. 2F for a representative experiment.

**Figure S3, related to Figure 3.**
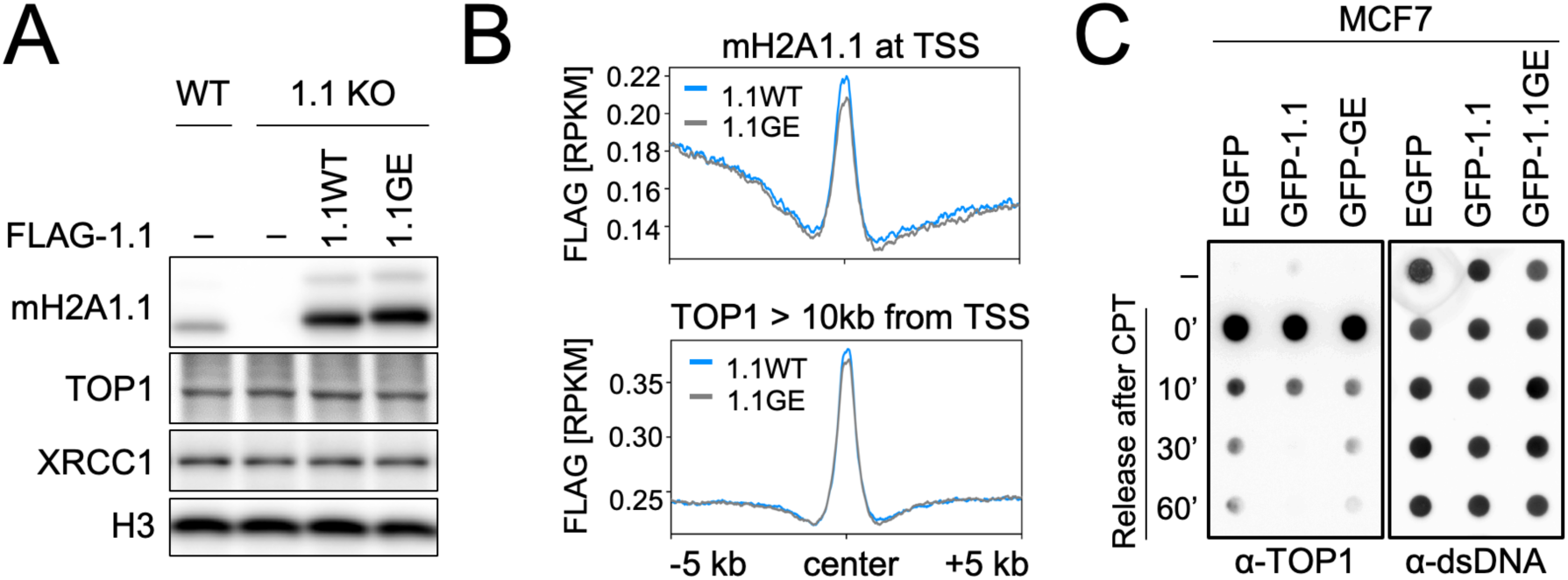
**(A)** Western blot for the indicated proteins in MDA-MB-231 WT or macroH2A1.1 knockout (1.1KO) cells, with or without stably integrated, FLAG-tagged wildtype (1.1WT) or G224E mutant macroH2A1.1 (1.1GE). **(B)** Profile plots for FLAG CUT&RUN signal in 1.1WT- or 1.1GE-expressing cells from (A). Signal was centered on TSS-proximal or TSS-distal TOP1 peaks. **(C)** RADAR assay as in Fig. 3A in MCF7 cells expressing GFP-tagged macroH2A1.1 (GFP-1.1) or macroH2A1.1 G224E (GFP-1.1GE), or an EGFP vector control.

**Figure S4, related to Figure 4.**
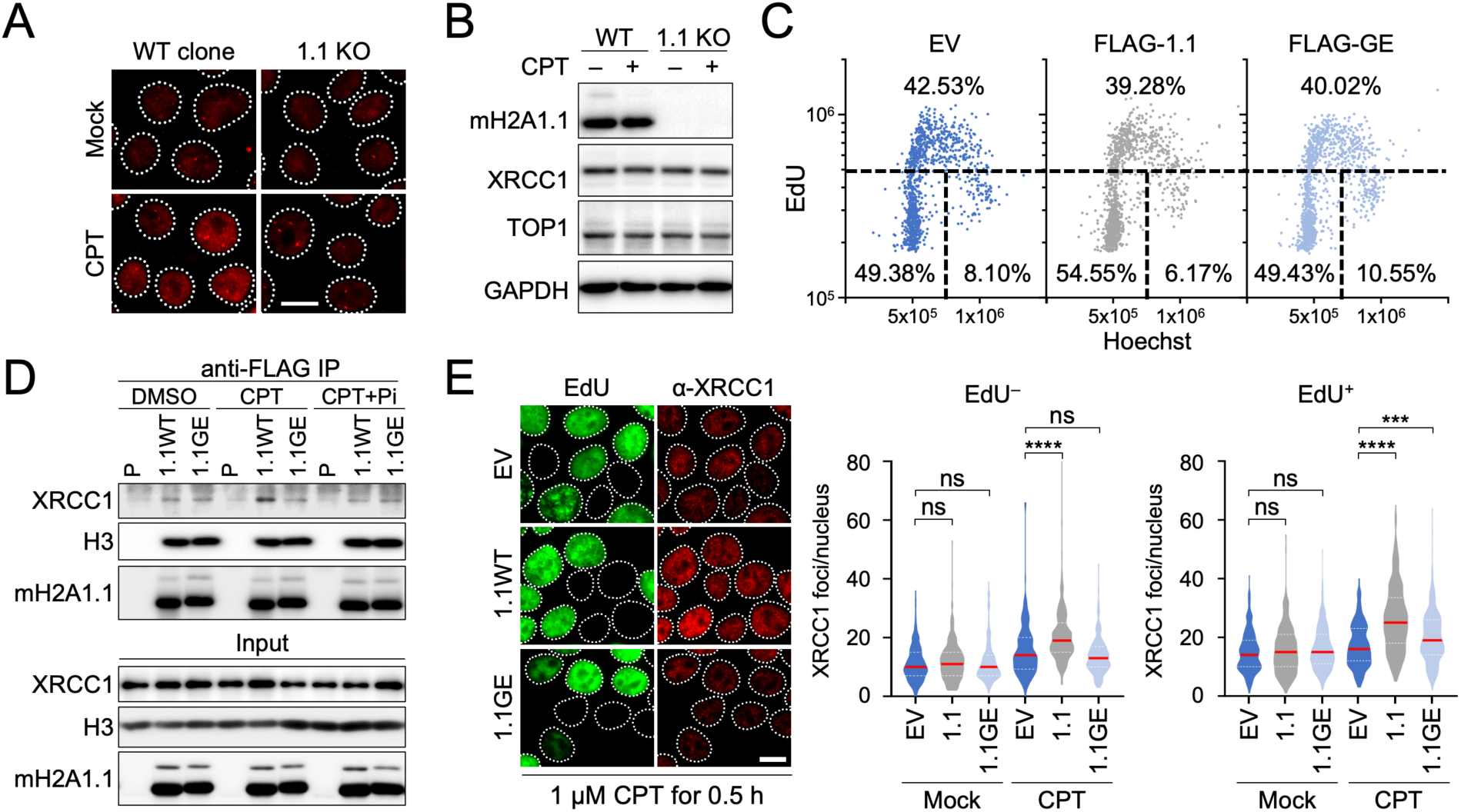
**(A)** Representative images of XRCC1 IF in MCF7 macroH2A1.1 knockout (1.1 KO) cells and a corresponding WT clone, quantified in Fig. 4B. Scale bar: 10 µm. **(B)** Western blot for the indicated proteins in cells from (A) in the presence or absence of 1 µM CPT for 30 min. **(C)** IF analysis of cell cycle profiles based on DNA content (Hoechst) and EdU incorporation in MCF7 cell lines from Fig. 4D, labeled with EdU for 30 min, EV: empty vector. **(D)** Western blot for the indicated proteins in nuclear lysates (input) or IP lysates from parental 293 cells (P) and cells stably expressing wild-type (WT) or G224E mutant FLAG-macroH2A1.1 (1.1GE) in the presence or absence of CPT and PARP inhibitor (Pi). **(E)** XRCC1 foci quantification and representative images in EdU^+^ and EdU^−^ cells from Fig. 4C; red lines reflect the median, P values are based on Mann-Whitney U test; *** p < 0.001, **** p < 0.0001, ns: not significant. Scale bar: 10 µm.

**Figure S5, related to Figure 5.**
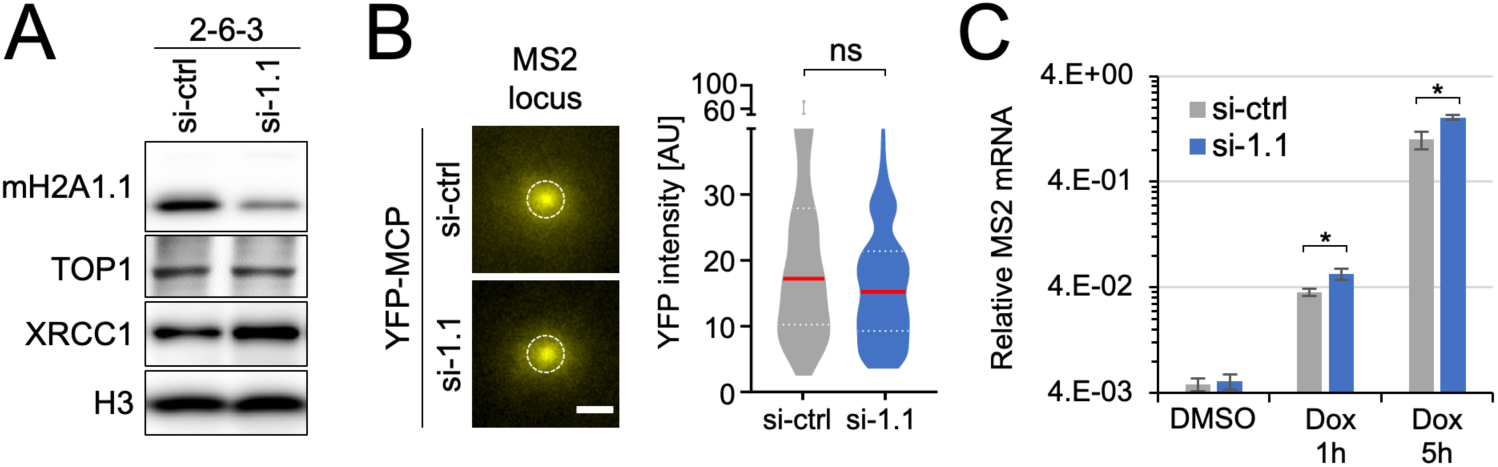
**(A)** Western blot for the indicated proteins in U2OS MS2-reporter cells expressing siRNA against macroH2A1.1 (si-1.1) or a non-targeting control siRNA (si-ctrl). **(B)** YFP-MCP intensity distribution after background subtraction in cells from Fig. 5B-C. Cells were transfected with a non-targeting control siRNA (si-ctrl) or siRNA against macroH2A1.1 (si-1.1) (n>70) and analyzed following 5 h Dox/PARGi treatment, scale bar: 1 µm. Red lines reflect the median, p value is based on Mann-Whitney U test, ns: not significant. **(C)** RT-PCR for MS2 transcript in cells expressing si-1.1 or si-ctrl treated with DMSO or Dox for the indicated time points, PARGi was added for 30 min. mRNA levels were normalized to β-actin and rpl13a housekeeping genes. Bar graphs depict mean and SD, p values are based on Student’s two-tailed t-test, * p < 0.05.

**Figure S6, related to Figures 6 and 7.**
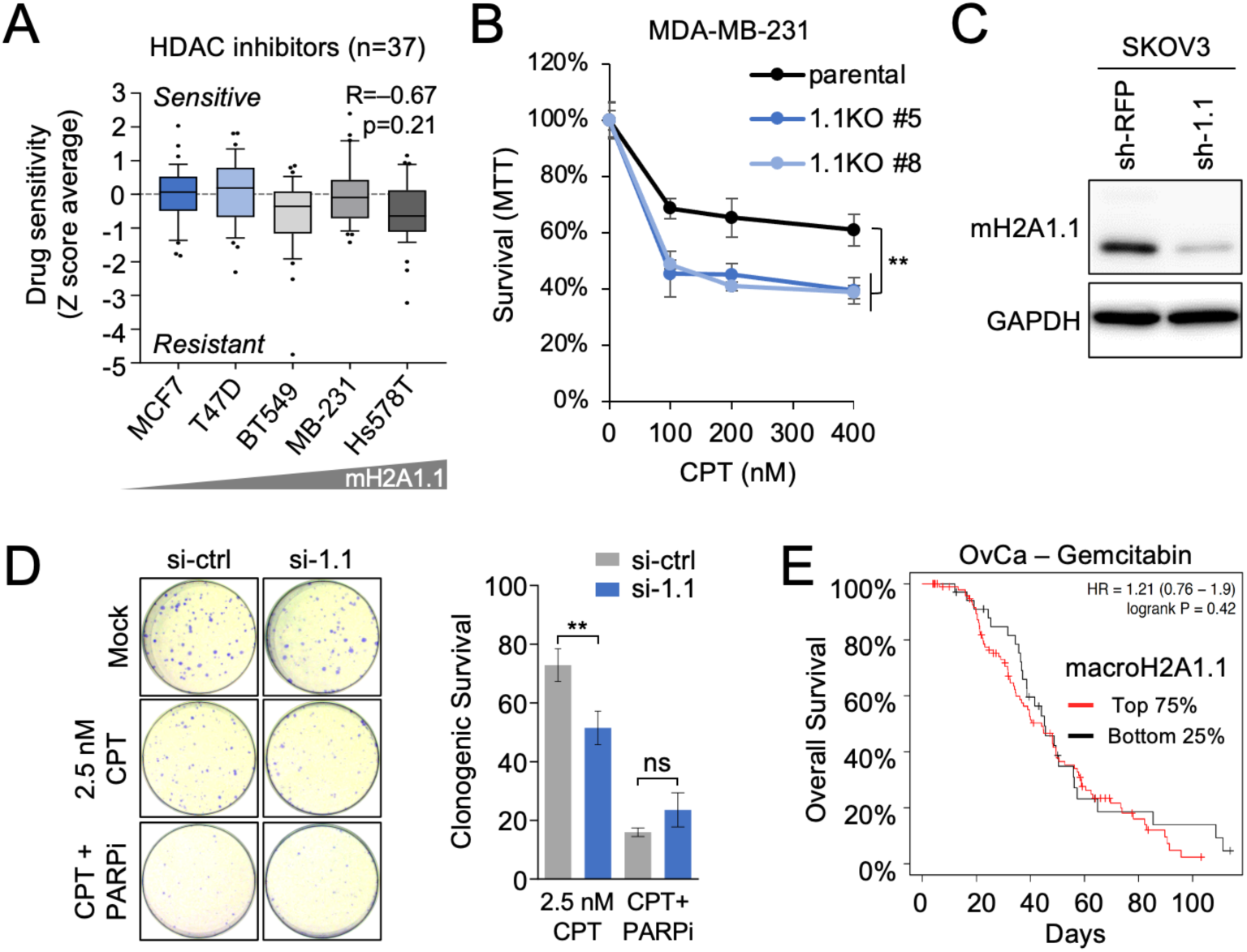
**(A)** HDAC inhibitor activity levels based on drug screen in NCI60 breast cancer cell lines, as in Fig. 6B; n=number of compounds tested, P value is based on Pearson’s Correlation Coefficient. **(B)** Cell viability of parental MDA-MB-231 cells and two independent macroH2A1.1 KO (1.1KO) knockout clones in response to CPT treatment, measured as in Fig. 6C, data are presented as mean and SD (n≥3). ** p < 0.01, based on Students two-tailed t-test, relative to parental MDA-MB-231 cells. **(C)** Western blot for the indicated proteins in SKOV3 cells stably expressing shRNAs against macroH2A1.1 or RFP (non-targeting control). **(D)** Clonogenic survival of SKOV3 ovarian cancer cells expressing the indicated siRNAs in response to the indicated drug combinations. Survival was normalized to untreated cells for each siRNA transfection. Representative images are shown, data are presented as mean and SD (n=3). ** p < 0.01, based on Student’s two-tailed t-test. **(E)** Kaplan-Meyer analysis of overall survival of TCGA ovarian cancer patient subgroups where treatment regimens contained gemcitabine (n=135). Patients were stratified by macroH2A1.1 mRNA expression as in Fig. 7B.

