## Supplemental Table 1 for "Epigenetic control of Topoisomerase 1 activity presents a cancer vulnerability"

**Supplemental Materials**

**Supplemental Table:**

**Table S1. Primers and small RNA sequences.**

| **Name** | **Sequence** | **Notes** |
| --- | --- | --- |
| **shRNA** |  |  |
| sh-1.1 | 5’-CGACAAACACTGACTTCTAC-3’ | This study |
| sh-1.2 | 5’ CTGAACCTTATTCACAGTGAA-3’ | Ref. 46 |
| sh-RFP | 5'-CGTAATGCAGAAGAAGACCAT-3' | Ref. 46 |
| **siRNA** |  |  |
| si-1.1 | 5’- CGACAAACACUGACUUCUA-3’ | Dharmacon |
| si-control | ON-TARGET Plus | Dharmacon |
| **gRNA** |  |  |
| E6b(1.1) KO 5’ | 5’-TTGGACCGAGACCCACGCAC-3’ | 1.1 exon-specific KO |
| E6b(1.1) KO 3’ | 5’ TTCTACATCGGTGGTGAAGT-3’ | 1.1 exon-specific KO |
| **Primers** |  |  |
| E6b KO-detect F | 5’- TGATCTCATGTGTGTGTTTCTCT-3’ | 1.1 KO screening |
| E6b KO-detect R | 5’-TGGCTGACAGCTAGCTTTATT-3’ | 1.1 KO screening |
| H3K27^low^ F | 5’-CCTTTCACCCAGTACCTCATTT-3’ | CUT&RUN validation |
| H3K27^low^ R | 5’-TCCTTCAATCCACCCATTCATC-3’ | CUT&RUN validation |
| H3K27^high^ F | 5’-GGTGGCTGTAACTCTCTCGT-3’ | CUT&RUN validation, Ref. 15 |
| H3K27^high^ R | 5’-CCAGGCCCCAGATGATAGAG-3’ | CUT&RUN validation, Ref. 15 |
| RPL13A RT F | 5’-GAAGTACCAGGCAGTGACAG-3’ | RT-PCR |
| RPL13A RT R | 5’-GGTCTTGAGGACCTCTGTG-3’ | RT-PCR |
| b-Actin RT F | 5’-TTCTACAATGAGCTGCGTGTGGCT-3’ | RT-PCR |
| b-Actin RT R | 5’-TCATCTTCTCGCGGTTGGCCT-3’ | RT-PCR |
| MS2-reporter F | 5’-TCATTAGATCCTGAGAACTTCA-3’ | RT-PCR, Ref. 36 |
| MS2-reporter R | 5’-TTTTGGCAGAGGGAAAAAGA-3’ | RT-PCR, Ref. 36 |

**Supplemental Figures**

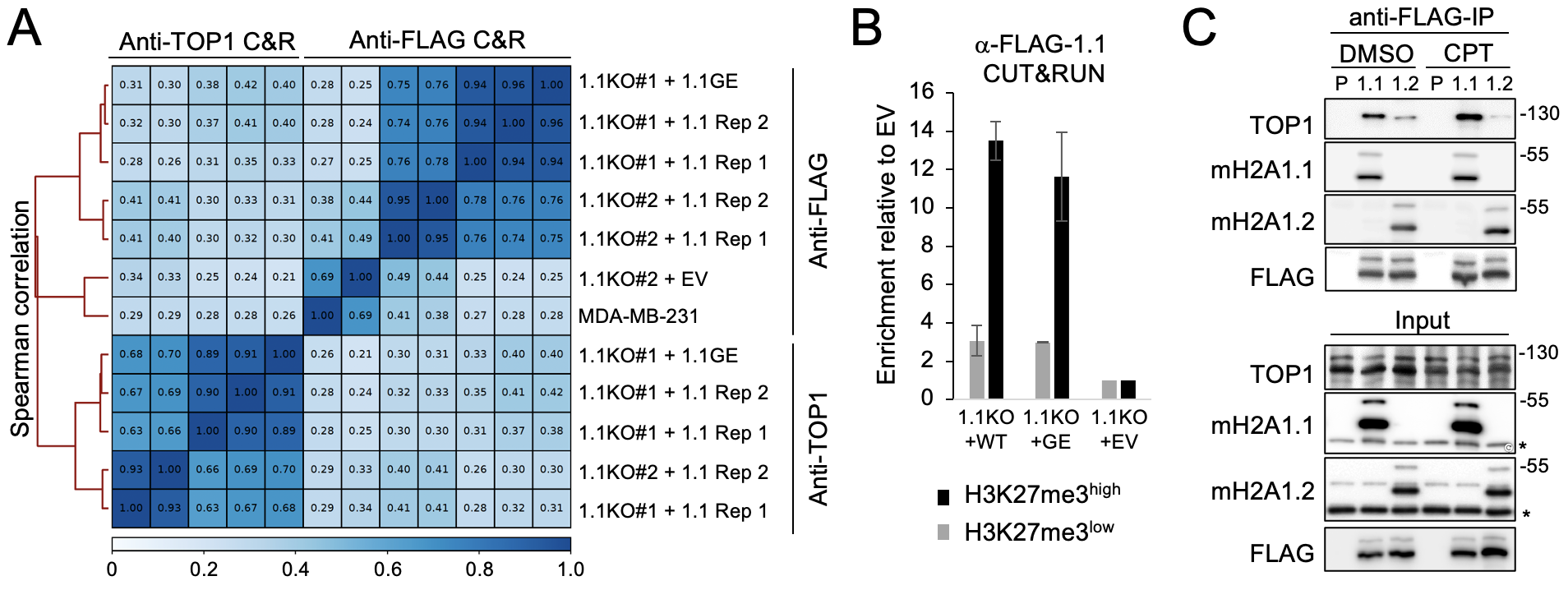

**Figure S1, related to Figure 1. (A)** Unsupervised hierarchical clustering of the indicated CUT&RUN NGS samples based on Spearman Correlation Coefficients. 1.1KO#1 and 1.1KO#2 represent independent CRISPR knockout clones, Rep 1 and Rep 2 independent experimental replicates. **(B)** qPCR analysis of FLAG-macroH2A1.1 CUT&RUN samples from MDA-MB-231 macroH2A1.1 knockout (1.1KO) cells reconstituted with WT or G224E (GE) mutant macroH2A1.1, or empty vector (EV) at an H3K27me3^high^ heterochromatin region known to bind macroH2A1.1 and an H3K27me3^low^ control region ^15^, see Table S1 for primer sequences. Samples were normalized to the EV control, values are expressed as mean and SD (n=2). **(C)** Western blot for the indicated proteins in nuclear lysates (input) or anti-FLAG IP samples from parental (P) 293 cells and FLAG-macroH2A1.1 (1.1) or FLAG-macroH2A1.2 (1.2) knock-in cells in the presence or absence of CPT treatment; * endogenous macroH2A1.1 or macroH2A1.2 protein.

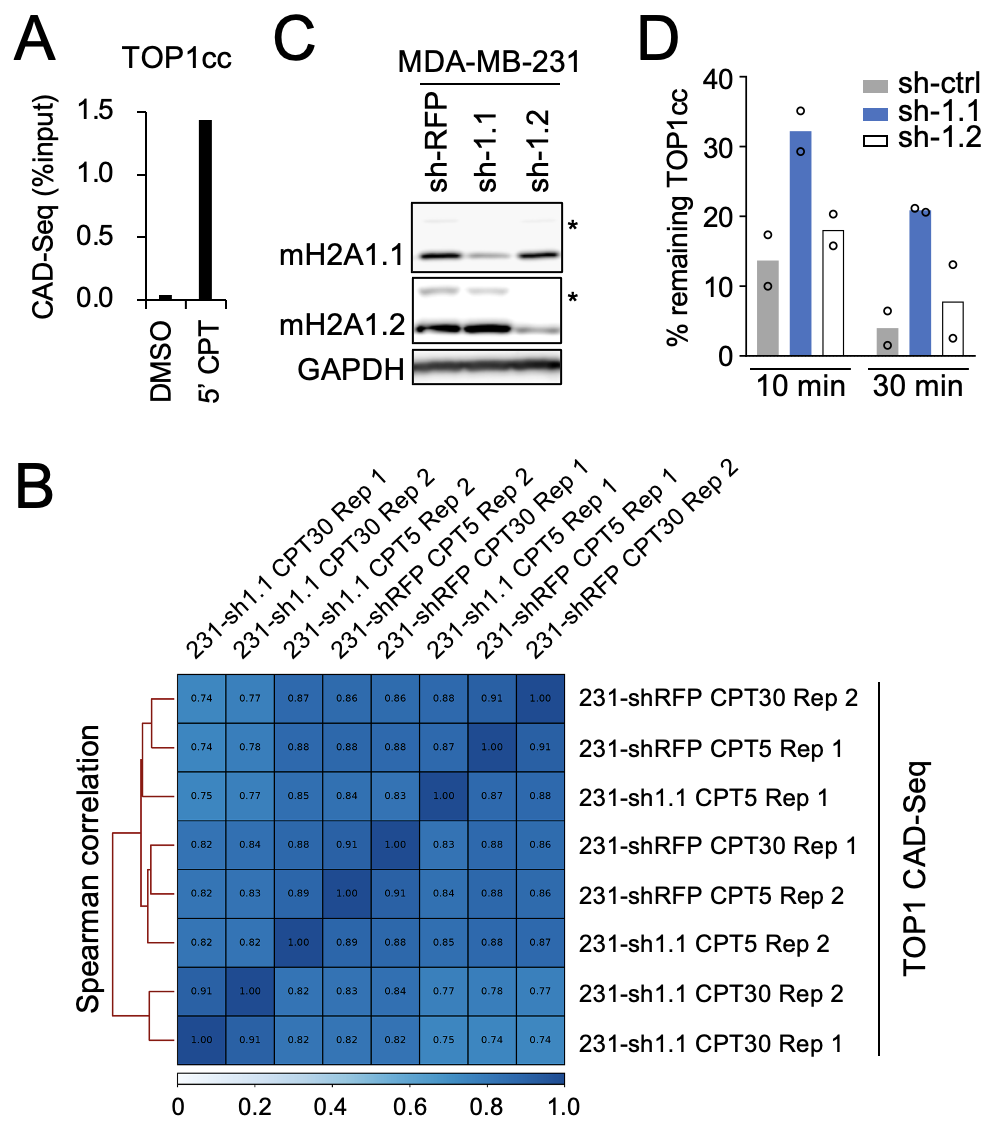

**Figure S2, related to Figure 2.** **(A)** qPCR analysis of CAD-Seq IP DNA relative to input at the TOP1cc-rich MYC promoter as described in Ref. 25, in MDA-MB-231 cells in the absence (DMSO) or presence of TOP1cc stabilization (CPT 5’). **(B)** Unsupervised hierarchical clustering of the indicated TOP1 CAD-Seq samples as in Fig S1A. CPT5 and CPT30: 5 min and 30 min CPT treatment, respectively; Rep: replicate experiment. **(C)** Western blot for the indicated proteins in MDA-MB-231 cells stably expressing shRNAs against macroH2A1.1, macroH2A1.2 or RFP (non-targeting control); * ubiquitinated macroH2A1 isoforms. **(D)** Quantification of TOP1cc turnover at the indicated time points after CPT treatment. The percent of remaining TOP1cc relative to 0’ after CPT is shown for two independent RADAR assays (open circles), see Fig. 2F for a representative experiment.

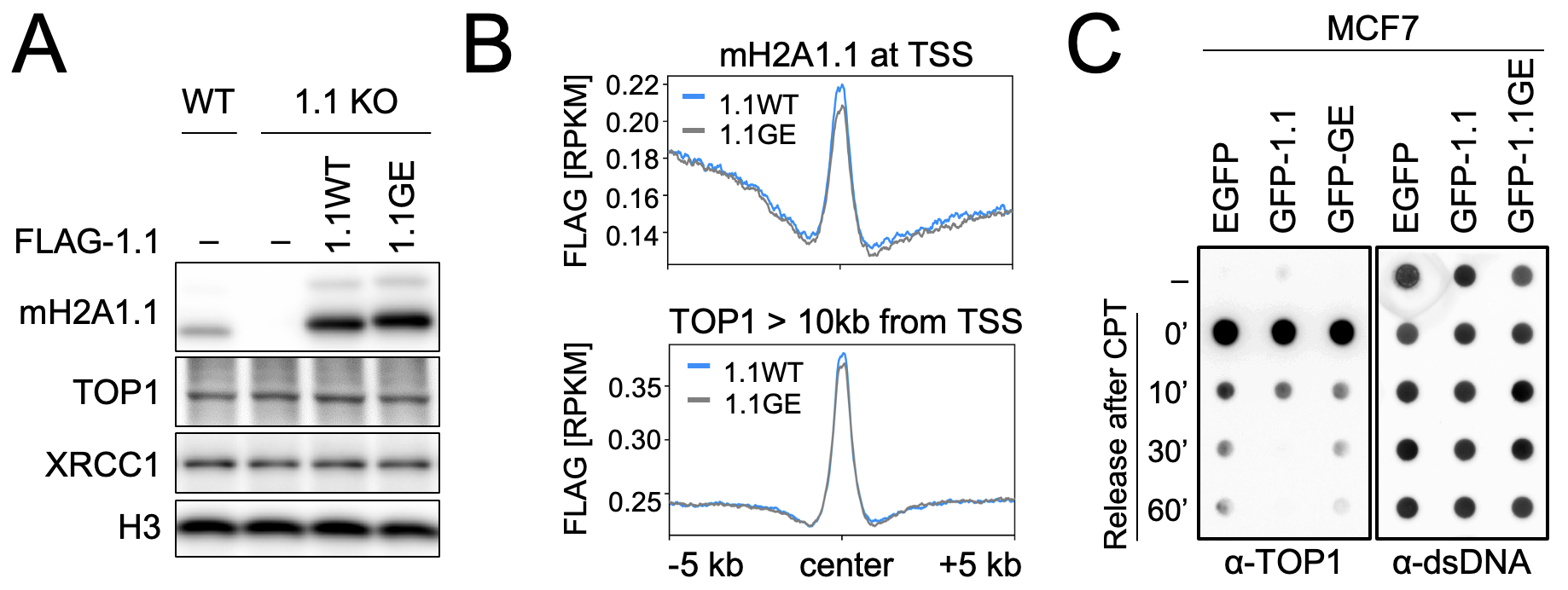

**Figure S3, related to Figure 3.** **(A)** Western blot for the indicated proteins in MDA-MB-231 WT or macroH2A1.1 knockout (1.1KO) cells, with or without stably integrated, FLAG-tagged wildtype (1.1WT) or G224E mutant macroH2A1.1 (1.1GE). **(B)** Profile plots for FLAG CUT&RUN signal in 1.1WT- or 1.1GE-expressing cells from (A). Signal was centered on TSS-proximal or TSS-distal TOP1 peaks. **(C)** RADAR assay as in Fig. 3A in MCF7 cells expressing GFP-tagged macroH2A1.1 (GFP-1.1) or macroH2A1.1 G224E (GFP-1.1GE), or an EGFP vector control.

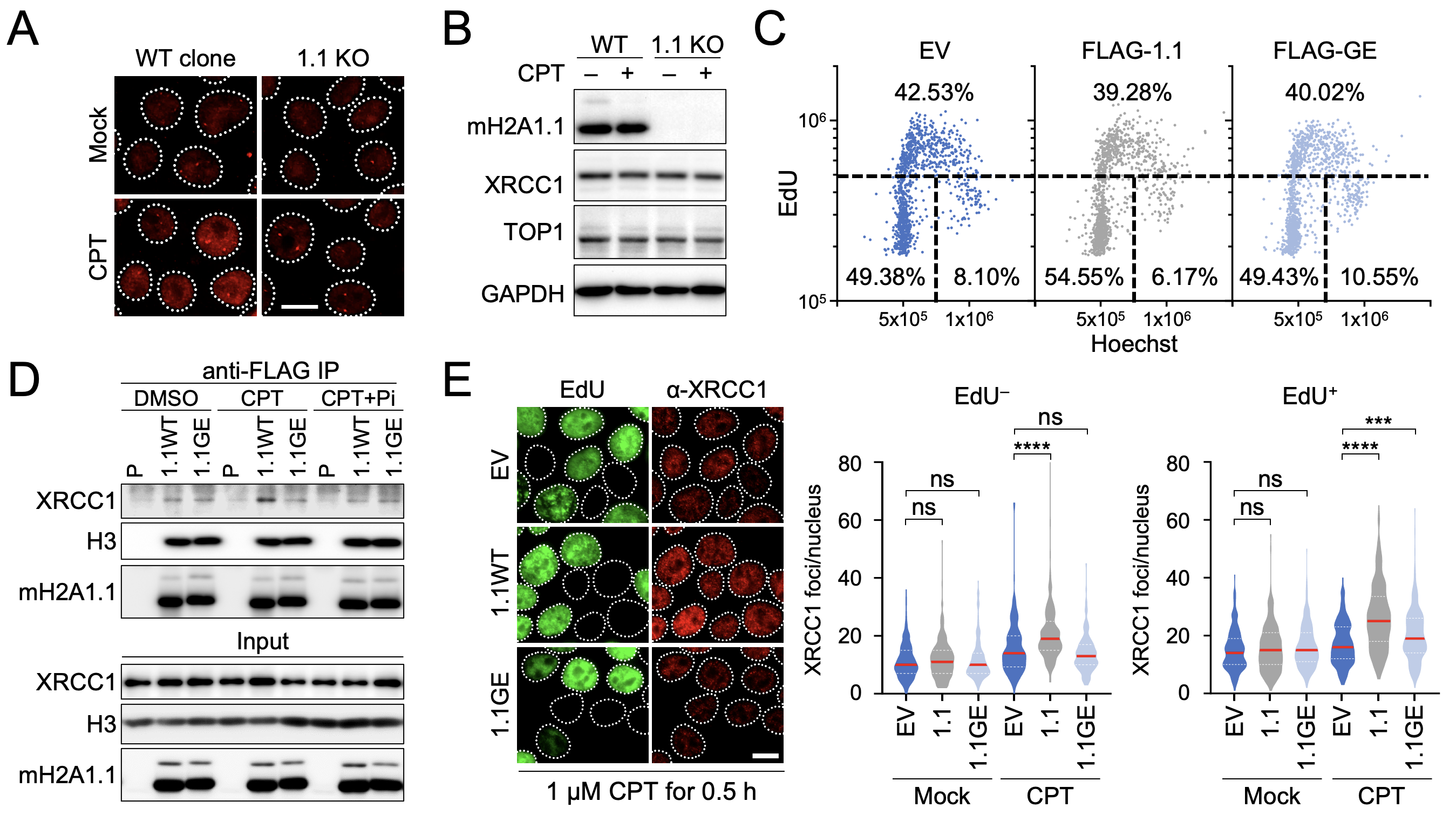

**Figure S4, related to Figure 4.** **(A)** Representative images of XRCC1 IF in MCF7 macroH2A1.1 knockout (1.1 KO) cells and a corresponding WT clone, quantified in Fig. 4B. Scale bar: 10 µm. **(B)** Western blot for the indicated proteins in cells from (A) in the presence or absence of 1 µM CPT for 30 min. **(C)** IF analysis of cell cycle profiles based on DNA content (Hoechst) and EdU incorporation in MCF7 cell lines from Fig. 4D, labeled with EdU for 30 min, EV: empty vector. **(D)** Western blot for the indicated proteins in nuclear lysates (input) or IP lysates from parental 293 cells (P) and cells stably expressing wild-type (WT) or G224E mutant FLAG-macroH2A1.1 (1.1GE) in the presence or absence of CPT and PARP inhibitor (Pi). **(E)** XRCC1 foci quantification and representative images in EdU^+^ and EdU^–^ cells from Fig. 4C; red lines reflect the median, P values are based on Mann-Whitney U test; *** p < 0.001, **** p < 0.0001, ns: not significant. Scale bar: 10 µm.

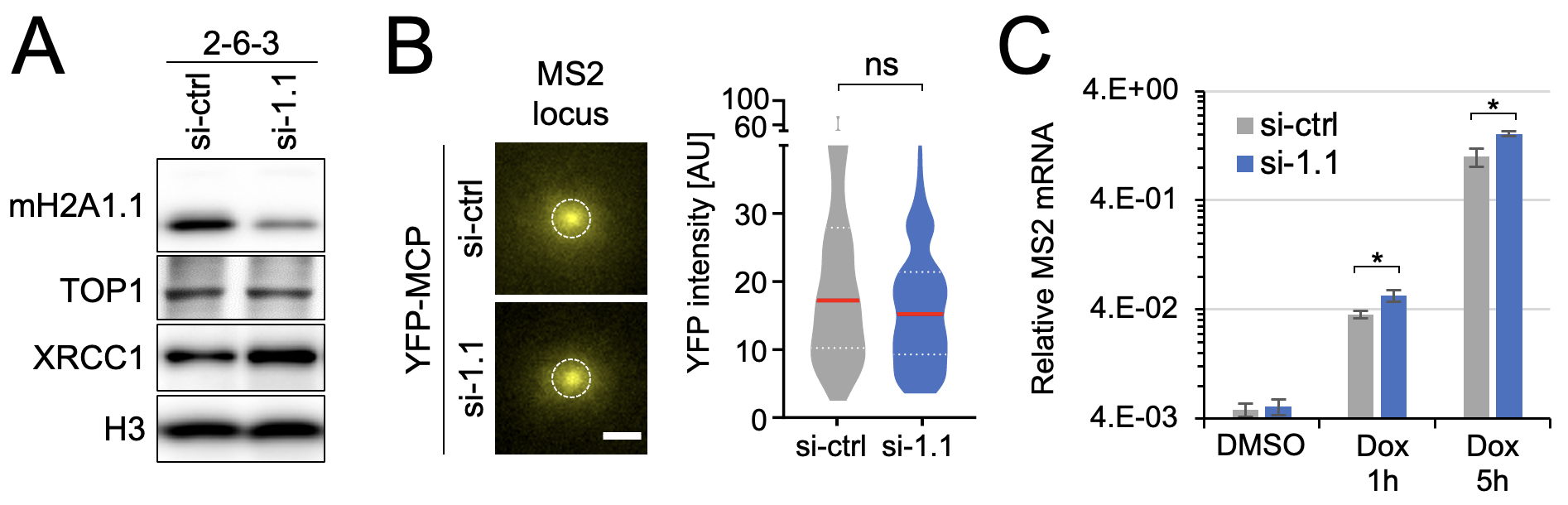

**Figure S5, related to Figure 5. (A)** Western blot for the indicated proteins in U2OS MS2-reporter cells expressing siRNA against macroH2A1.1 (si-1.1) or a non-targeting control siRNA (si-ctrl). **(B)** YFP-MCP intensity distribution after background subtraction in cells from Fig. 5B-C. Cells were transfected with a non-targeting control siRNA (si-ctrl) or siRNA against macroH2A1.1 (si-1.1) (n>70) and analyzed following 5 h Dox/PARGi treatment, scale bar: 1 µm. Red lines reflect the median, p value is based on Mann-Whitney U test, ns: not significant. **(C)** RT-PCR for MS2 transcript in cells expressing si-1.1 or si-ctrl treated with DMSO or Dox for the indicated time points, PARGi was added for 30 min. mRNA levels were normalized to β-actin and rpl13a housekeeping genes. Bar graphs depict mean and SD, p values are based on Student’s two-tailed t-test, * p < 0.05.

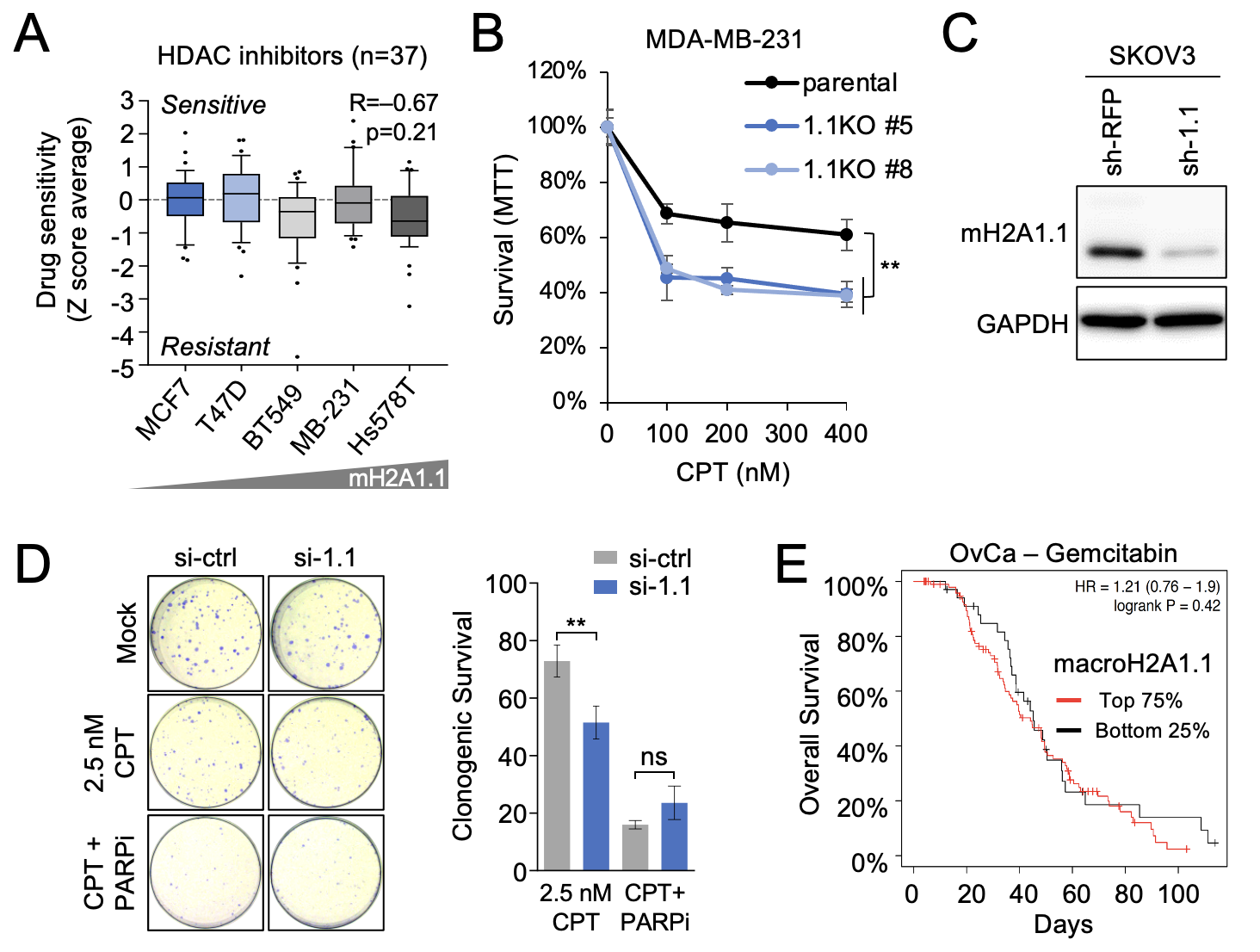

**Figure S6, related to Figures 6 and 7.** **(A)** HDAC inhibitor activity levels based on drug screen in NCI60 breast cancer cell lines, as in Fig. 6B; n=number of compounds tested, P value is based on Pearson’s Correlation Coefficient. **(B)** Cell viability of parental MDA-MB-231 cells and two independent macroH2A1.1 KO (1.1KO) knockout clones in response to CPT treatment, measured as in Fig. 6C, data are presented as mean and SD (n≥3). ** p < 0.01, based on Students two-tailed t-test, relative to parental MDA-MB-231 cells. **(C)** Western blot for the indicated proteins in SKOV3 cells stably expressing shRNAs against macroH2A1.1 or RFP (non-targeting control). **(D)** Clonogenic survival of SKOV3 ovarian cancer cells expressing the indicated siRNAs in response to the indicated drug combinations. Survival was normalized to untreated cells for each siRNA transfection. Representative images are shown, data are presented as mean and SD (n=3). ** p < 0.01, based on Student’s two-tailed t-test. **(E)** Kaplan-Meyer analysis of overall survival of TCGA ovarian cancer patient subgroups where treatment regimens contained gemcitabine (n=135). Patients were stratified by macroH2A1.1 mRNA expression as in Fig. 7B.
